# Morphology of obligate ectosymbionts reveals *Paralaxus* gen. nov., a new circumtropical genus of marine stilbonematine nematodes

**DOI:** 10.1101/728105

**Authors:** Florian Scharhauser, Judith Zimmermann, Jörg A. Ott, Nikolaus Leisch, Harald Gruber-Vodicka

**Author notes:** Contributed equally (Florian Scharhauser, Judith Zimmermann). Harald Gruber-Vodicka, Max Planck Institute for Marine Microbiology, Celsiusstrasse 1; 28359 Bremen, Germany.

## Abstract

Stilbonematinae are a subfamily of conspicuous marine nematodes, distinguished by a coat of sulphur-oxidizing bacterial ectosymbionts on their cuticle. As most nematodes, the worm hosts have a simple anatomy and few taxonomically informative characters, and this has resulted in numerous taxonomic reassignments and synonymizations. Recent studies using a combination of morphological and molecular traits have helped to improve the taxonomy of Stilbonematinae but also raised questions on the validity of several genera. Here we describe a new circumtropically distributed genus *Paralaxus* (Stilbonematinae) with three species: Paralaxus cocos sp. nov., *P. bermudensis* sp. nov. and *P. columbae* sp. nov.. We used single worm metagenomes to generate host 18S rRNA and cytochrome oxidase I (COI) as well as symbiont 16S rRNA gene sequences. Intriguingly, COI alignments and primer matching analyses suggest that the COI is not suitxable for PCR-based barcoding approaches in Stilbonematinae as the genera have a highly diverse base composition and no conserved primer sites. The phylogenetic analyses of all three gene sets however confirm the morphological assignments and support the erection of the new genus *Paralaxus* as well as corroborate the status of the other stilbonematine genera. *Paralaxus* most closely resembles the stilbonematine genus *Laxus* in overlapping sets of diagnostic features but can be distinguished from *Laxus* by the morphology of the genus-specific symbiont coat. Our re-analyses of key parameters of the symbiont coat morphology as character for all Stilbonematinae genera show that with amended descriptions, including the coat, highly reliable genus assignments can be obtained.

## Introduction

The identification of many nematode genera and species is difficult based on morphological characters alone (Sudhaus & Kiontke 2007, de-León & Nadler 2010, Derycke et al. 2008, Palomares-Rius et al. 2014). A prime example for this problem are the *Stilbonematinae*, a subfamily of the *Desmodoridae* that have experienced several changes in the classification of species and even genera in the past (Tchesunov 2013; Ott et al. 2014). The exclusively marine *Stilbonematinae* are common members of the interstitial meiofauna in sheltered intertidal and subtidal porous sediments and have been found worldwide (Ott et al. 2004, Tchesunov et al. 2013). Their diversity and abundances are highest in subtropical and tropical shallow-water sands, but some species were also described from higher latitudes (e.g. Gerlach 1950, Wieser 1960, Platt & Zhang 1982, Riemann et al. 2003, Tchesunov 2012), deep-sea sediments (Van Gaever et al. 2004, Tchesunov et al. 2012, Leduc 2013), near shallow hydrothermal vents (Kamenev et al. 1993, Thiermann et al. 1997) and methane seeps (Dando et al. 1995).

To date, a total of 10 different stilbonematine nematode genera and more than 50 different species are described (reviewed by Tchesunov 2013; Leduc 2013, Armenteros et al. 2014b, Ott et al. 2014a,b, Leduc and Sinniger 2018). Most species and genera still lack molecular data, but recent taxonomic work has started to integrate molecular data using the 18S rRNA gene (18S) (Ott et al. 2014a, b; Armenteros et al. 2014ab; Leduc and Zhao 2016, Leduc and Sinniger 2018) and the mitochondrial cytochrome oxidase subunit I (COI) gene (Armenteros et al. 2014a, b). The phylogenetic signal of the two marker genes was however not consistent and called morphological genus and species identifications into question (Armenteros et al. 2014b). While the monophyly of the subfamily *Stilbonematinae* has strong support based on 18S data (Kampfer et al. 1998, Bayer et al. 2009, van Megen et al. 2009, Ott et al 2014a and b; Leduc & Zhao 2016), it was questioned in a recent study by Armenteros et al. (2014b) which used both 18S and COI.

A trait that unites all stilbonematine nematodes is the conspicuous ectosymbiosis with sulphur-oxidizing *Gammaproteobacteria* that cover major parts of the cuticle (reviewed by Ott et al. 2004). Stilbonematine symbionts have diverse morphologies, including rod, coccus, coccobacillus, filament, corn-kernel or crescent shapes, and each host species harbours a single morphotype (Ott & Novak 1989, Polz et al. 1992; Polz et al. 1994; Ott et al., 2004; Bayer et al. 2009, Pende et al 2014, Ott et al. 2014a, b). In phylogenetic analyses of the symbiont 16S rRNA gene (16S) sequences, all stilbonematine symbionts fall into a monophyletic clade of *Gammaproteobacteria* related to *Chromatiaceae*, which has been described as *Candidatus* Thiosymbion (from here on ‘Thiosymbion’) (Bayer et al. 2009, Bulgheresi et al. 2011, Heindl et al. 2011, Zimmermann et al. 2016). Based on phylogenetic analyses of host 18S and symbiont 16S sequences closely related stilbonematine nematode species of the same genus consistently harbour closely related and genus specific ‘Thiosymbion’ symbionts (Ott et al. 2004, Zimmermann et al. 2016). ‘Thiosymbion’ encompasses not only the ectosymbionts of stilbonematine nematodes, but also the primary symbionts of gutless phallodriline oligochaetes and siphonolaimid nematodes (Zimmermann et al. 2016). Comparative analyses of stilbonematine nematodes and their symbionts showed a high degree of phylogenetic congruence that indicates a long-term co-diversification between ‘Thiosymbion’ and the *Stilbonematinae* (Zimmermann et al. 2016).

Recent 18S-based phylogenetic analyses suggested a novel, morphologically uncharacterized stilbonematine genus from the Caribbean Sea (Belize) and the Australian Great Barrier Reef (Heron Island) (Zimmermann et al. 2016). The genus level of this clade was supported by the distinct 16S sequences of their symbionts that also formed a new phylogenetic clade in the study. Here we combine morphological and molecular data to describe this recently detected host clade as ‘*Paralaxus*’ gen. nov.. We use single worm metagenomes to provide host 18S and COI gene sequences as well as symbiont 16S sequences from nine *Paralaxus* specimens, and 20 additional specimens covering six stilbonematine genera. We place *Paralaxus* in a unified taxonomy of *Stilbonematinae* and formally describe the new genus *Paralaxus* with a Caribbean type species *P. cocos* sp. nov. In addition, we describe two new species, *P. bermudensis* sp. nov. and *P. columbae* sp. nov. from Bermuda and Florida, and report on members of this new genus from localities in the Western and Central Pacific Ocean. While characterizing the key anatomical features of *Paralaxus*, we identified the high taxonomic value of its symbiont coat. We extend this concept and evaluate coat and symbiont morphology as additional traits for all stilbonematine genera.

In addition, we substantially improve the molecular dataset of *Stilbonematinae* full-length 18S and COI marker gene sequences to answer the following questions: (1) Are *Stilbonematinae* monophyletic? (2) At which taxonomic level are 18S and COI valuable phylogenetic markers for *Stilbonematinae*? (3) Can we find suitable COI PCR priming sites that are conserved across the subfamily?

## Material and Methods

### Meiofauna collection, preparation and microscopy

Sediment samples were collected in the Caribbean Sea (Belize), the Western Atlantic Ocean (Florida and Bermuda), the Western and Central Pacific Ocean (Australia and Hawaii) as well as the North and Mediterranean Sea (Fig. 1 and S1) using cores or buckets. Nematodes were extracted by decantation through a 64 µm mesh sieve and sorted live under a dissecting microscope. Specimens were fixed in 4% formaldehyde for taxonomic preparations or in 2.5% glutaraldehyde in 0.1 M sodium cacodylate buffer and were post fixed in 2% OsO_4_ (for scanning electron microscopy, SEM) and stored at 4 °C until further analyses. Before fixation, live specimens for sequencing were photographically vouchered in the field *i.e*. micrographs were taken of the full specimen mounted in seawater, capturing the head region with mouth, amphids and pharynx, the anal and tail region, the symbiont coat, and male and female reproductive organs. After imaging, specimens were washed in 0.2 µm filtered seawater and immediately fixed in either RNAlater® stabilization solution (Ambion, Foster City, US), 70% ethanol or pure methanol and stored at 4° C until the molecular analyses.

**Fig. 1.**
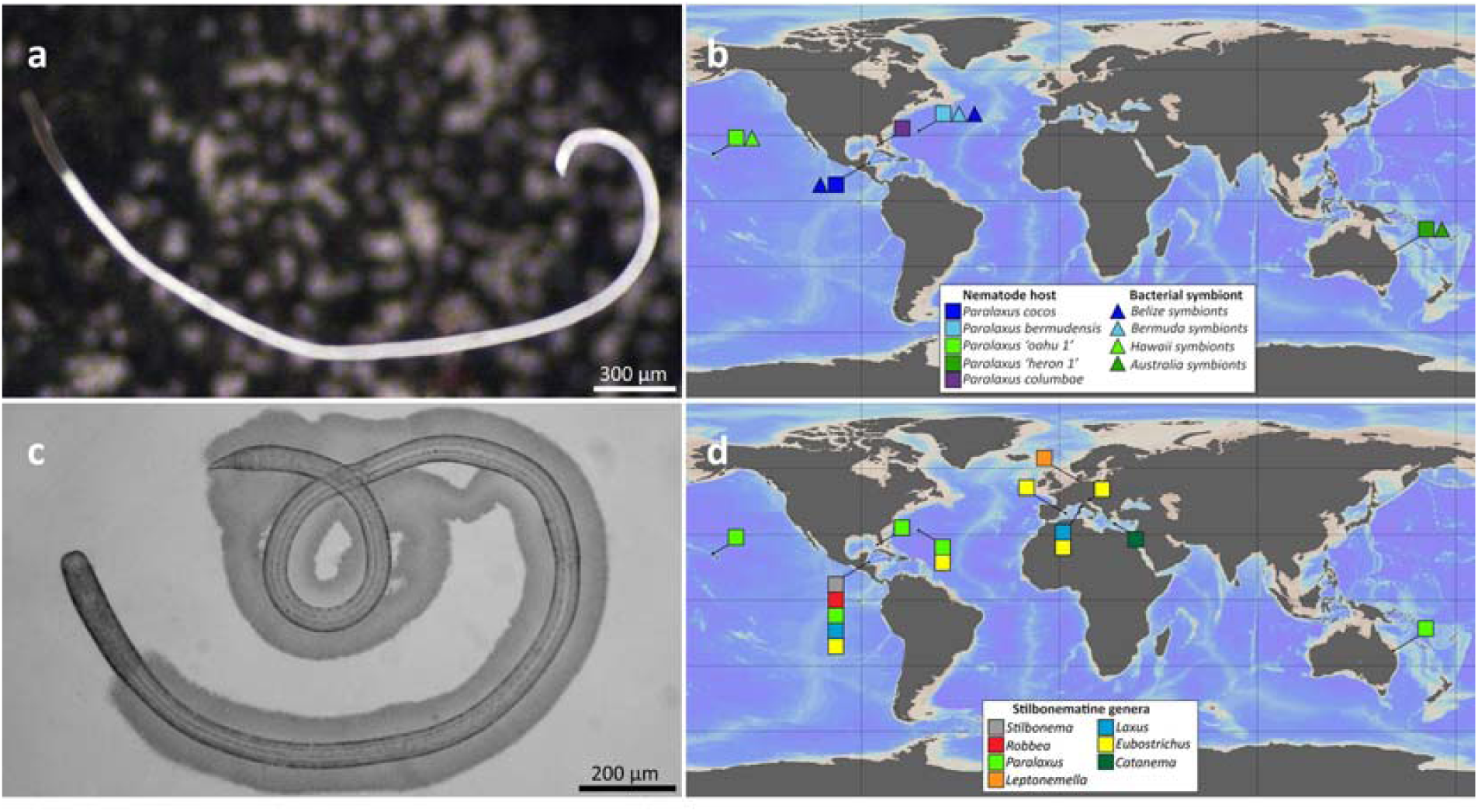
*Paralaxus* gen. nov. habitus and sampling sites of stilbonematine nematodes. (a) Micrograph of Paralaxus *bermudensis* sp. nov. The symbiont free anterior region is clearly visible compared to the densely coated white body. (b) Sampling locations of *Paralaxus* species included in this study (c) Juvenile specimen of *Paralaxus columbae* sp. nov. with thick symbiotic coat and bacteria free head region. (d) Sampling locations of other stilbonematine nematode species included in this study. Specimen for this study were sampled from shallow-water sandy sediments in the Caribbean Sea, the Atlantic and the Pacific Ocean as well as the North and Mediterranean Sea. For details see Table S1 (Supporting information).

For detailed light microscopy, the formaldehyde or glutaraldehyde-fixed specimens were transferred into glycerol: water 1:9, slowly evaporated and finally mounted in pure glycerol. Drawings were made using a camera lucida on a Diavar microscope (Reichert, Vienna, Austria). Nomarsky interference contrast photos were taken on a Polyvar (Reichert, Vienna, Austria). Specimens for SEM were critical point dried and coated with palladium using a JEOL JFC-2300HR sputter coater (JEOL Ltd., Akishima, Japan) and examined on a JEOL IT 300 (JEOL Ltd., Akishima, Japan) at high vacuum mode.

### DNA extraction, marker gene amplification and metagenome sequencing

DNA from single nematodes including their attached ectosymbionts was extracted with the DNAeasy Blood and Tissue Micro Kit (Qiagen, Hilden, Germany) following manufacturer’s instructions with two amendments to the protocol – the proteinase K digestion was extended to at least 24 h and the elution step was performed twice with 20 µl of elution buffer. DNA concentrations were quantified using a Qubit 2.0 and the high sensitivity assay (Life Technologies, Carlsbad, US).

To characterize nematode hosts prior to metagenome sequencing we used both the ribosomal 18S and mitochondrial COI marker genes for PCR-based screenings. To amplify an approximately 600 bp long fragment of the nematode 18S we designed a nematode-specific primer pair: 3FNem 5’-GTTCGACTCCGGAGAGGGA-3’ and 5R 5’- CTTGGCAAATGCTTTCGC-3’. Amplification of the mitochondrial COI gene of several stilbonematine nematode genera including *Paralaxus* was conducted using the published primer pair 1490F 5’-GGTCAACAAATCATAAAGATATTGG-3’ and 2198R 5’- TAAACTTCAGGGTGACCAAAAATCA-3’ (Folmer et al. 1994). For 18S PCRs we used the Phusion® DNA polymerase, HF buffer (Finnzymes, Finland) and the following cycling conditions: 95° C for 5 min, followed by 38 cycles of 95° C for 1 min, 55° C for 1.5 min, 72° C for 2 min and followed by 72° C for 10 min. COI PCRs were conducted using the TaKaRa Taq™ DNA polymerase, 10x reaction buffer (Takara Bio Inc., Japan) and the following cycling conditions: 5 min at 95° C, then 36 cycles of 1 min at 95°C, 1.5 min at 42° C, and 2 min at 72° C, followed by 72° C for 10 min. While the 18S amplifications were successful for all nematode specimens, we could not obtain a single PCR product for the COI amplifications of any stilbonematine nematode specimen. In contrast, DNA extracts from *Lamellibrachia* tubeworms (Siboglinidae, Annelida) that were used as positive control always resulted in COI PCR products of the expected size.

We performed single specimen shotgun metagenome sequencing to generate data from the host nuclear genome, host mitochondria and symbionts using Illumina short-read technology. For Illumina sequencing 1 - 5 ng DNA per specimen were used for an Ovation Ultralow Library Systems kit (NuGEN Technologies, San Carlos, US). Illumina library preparation including size-selection on an agarose gel was performed at the Max Planck Genome Centre in Cologne, Germany. 2⍰× ⍰100 bp paired-end reads were sequenced on an Illumina HiSeq3000 (San Diego, US) for each individual (8 - 25 million reads each). We intended to extract the triple gene sets (symbiont 16S, host 18S and COI) for all specimens but differing assembly qualitites due to the very low DNA input did not allow us to retrieve all marker genes for each worm (Table S1). Host 18S and symbiont 16S gene sequences were generated from the raw reads with phyloFlash (Gruber-Vodicka et al. 2017, https://github.com/HRGV/phyloFlash). The COI was assembled using the following approach: In brief, after removing adapters and low-quality bases (Q < 2) with bbduk (Bushnell 2014), we retained all reads with a minimal length of 36 bp after quality trimming and conducted combined host and symbiont draft genome assemblies for all samples with SPAdes 3.1 – 3.9 (Bankevich et al. 2012. The contig containing the COI gene was then extracted from each assembly using BLAST 2.26+ searches with the available Stilbonematinae COI sequences as implemented in the Geneious software v. 11.1 (54) (Biomatters, New Zealand). Full-length COI genes were then predicted from the contigs using the Geneious gene prediction tool.

### COI compositional analyses and primer matching

All compositional and primer matching analyses were conducted with the 27 full-length stilbonematine nematode COI sequences we derived from the metagenomics data. Nucleotide composition was analyzed in Geneious. A primer mismatch analysis of the two COI primer sets: 1490F: 5’-GGT CAA CAA ATC ATA AGA TAT TGG-3’ and 2198R: 5’-TAA ACT TCA GGG TGA CCA AAA AAT CA-3’ (Folmer et al. 1994) and JB3F: 5’-TTT TTT GGG CAT CCT GAG GTT TAT-3’ and JB5 R: 5’-AGC ACC TAA ACT TAA AAC ATA ATG AAA ATG-3’ used by Armenteros et al. (2014 a,b), was performed using the MOTIF search option integrated in Geneious, with a maximum mismatch rate range from 1 - 7 for every primer. The mismatches were mapped on a MAFFT v.7 (Katoh & Standley 2013) alignment. The design of a new stilbonematine-specific COI primer set was conducted with Primer 3 (Untergasser et al. 2012). In brief, the consensus sequence of our full-length COI MAFFT alignment (1 bp to 1577 bp) was screened for a stilbonematine nematodes-specific primer set by applying Tm calculation after SantaLucia (1998) and the following parameters: primer size range 18 to 27 bp, melting temperature (Tm) range 57° C to 63° C and a GC percentage range 20 to 80. In addition, separate searches were conducted for the regions 1 bp to 750 bp and 750 bp to 1577 bp to facilitate the detection of primer pairs that cover similar ranges as the existing primer sets that only amplify parts of the gene.

### Phylogenetic analyses of host and symbiont genes

For the host 18S-based phylogenetic reconstruction we used 23 full-length sequences extracted from the single worm metagenomes together with 55 previously published and non-chimeric stilbonematine nematode 18S sequences longer than 1300 bp available in GenBank. All published non-stilbonematine nematode sequences of the order *Desmodorida* and *Chromadorida* sequences longer than 1300 bp available in 10/2018 served as outgroup.

In an extended 18S dataset with a total of 250 sequences, we included eight partial 18S sequences of five stilbonematine species from Armenteros et al. (2014b) and all non-chimeric 18S sequences of *Desmodorida* and *Chromadorida* with a minimum length of 700 bp available in GenBank.

For the host COI phylogeny, we constructed a COI matrix using the 27 full-length stilbonematine COI sequences we extracted from the metagenomics data of nine *Paralaxus* specimens, together with sequences of 18 specimens from six different stilbonematine genera (*Catanema, Eubostrichus, Laxus, Leptonemellla, Robbea and Stilbonema*). We added the full-length COI sequence from *Baylisascaris procyonis* (JF951366) as outgroup. All 28 nucleotide COI sequences were translated into protein sequences in Geneious using the invertebrate mitochondrial translation Table 5. In an extended COI matrix, we included partial COI sequences of the five species from Armenteros et al. (2014b) that were also included in the extended 18S dataset. Due to the high morphological similarity between *Paralaxus cocos* sp. nov. and *Leptonemella brevipharynx* (Armenteros et al. 2014b) the partial COI sequence of *L. brevipharynx* was also included in the analysis.

For the symbiont phylogenetic reconstruction, we used the 21 new 16S sequences that we extracted from the metagenomes as well as the 54 ‘Thiosymbion’ 16S genes sequences available in GenBank. We also included sequences of closely related uncultured *Gammaproteobacteria* (JF344100, JF344607, JF344324), and used five cultured free-living gammaproteobacterial sulfur-oxidizing bacteria from the Chromatiaceae (*Allochromatium vinosum* strain DSM180, *Marichromatium purpuratum* 984, *Thiorhodococcus drewsii* AZ1, *Thiocapsa marina* 5811, *Thiorhodovibrio* sp. 970) as an outgroup.

The host and symbiont ribosomal rRNA sequences were independently aligned using MAFFT v7 (Katoh & Standley 2013) with the Q-INS-I mode (Katoh & Toh 2008) as it considers the predicted secondary structure of the RNA. All COI sequences were translated into AA sequences and aligned in MAFFT v7 with the E-INS-i mode. The optimal substitution model for each alignment was assessed using ModelFinder (Kalyaanamoorthy et al., 2017). The mtZOA+F+G4 model was the best fit for the COI alignment, TIM3e+I+G4 for the 18S alignment and TIM+F+I+G4 for the 16S alignment. Phylogenetic trees were reconstructed using the maximum likelihood-based software IQTREE (Nguyen et al. 2015) utilizing the Ultrafast Bootstrap Approximation UFBoot (Minh et al., 2013) to assess node stability (10000 bootstrap runs). In addition to maximum likelihood, support values were generated using approximate Bayes (aBayes) (Anisimova et al. 2011) and SH-aLRT analyses (Guindon et al., 2010).

### Data accessibility

Sequences from this study were submitted to the European Nucleotide Archive (ENA) via GFBio data submission (Diepenbroek at el. 2014; https://www.gfbio.org) and are available under the accession number PRJEB27096.

## Results

### Descriptions

We analyzed 33 *Paralaxus* gen. nov. specimens from seven locations in the Pacific, the Atlantic and the Caribbean using light- and electron microscopy as well as molecular analyses based on marker genes extracted from single specimen metagenomes.

#### *Paralaxus* gen. nov

ZooBank registration (http://zoobank.org): urn:lsid:zoobank.org:act:417691DB-5EE3-49B7-B838-1BBF1569235F

GenBank entries for 18S rRNA and COI sequences: see Table S1.

We erect a new genus for this clade with the following characters: cephalic capsule without block layer (Fig. 2b, e, i, l, m); amphidial fovea spiral, no sexual dimorphism in shape of fovea (Fig. 2b, g, i, l, m); male tail with velum (Fig. 2c, j, n); symbiotic bacteria coccobacilli, arranged as multi-layered coat covering body except for anterior and posterior end (Fig. 1a, b). All *Paralaxus* sequences form statistically supported cIades in phylogenetic analyses of the 18S as well as the COI datasets (Fig. 3a and b). The specimens for each Paralaxus species clustered together in both gene sets and clearly separated each *Paralaxus* species from each other (Fig. 3a and b).

**Fig. 2.**
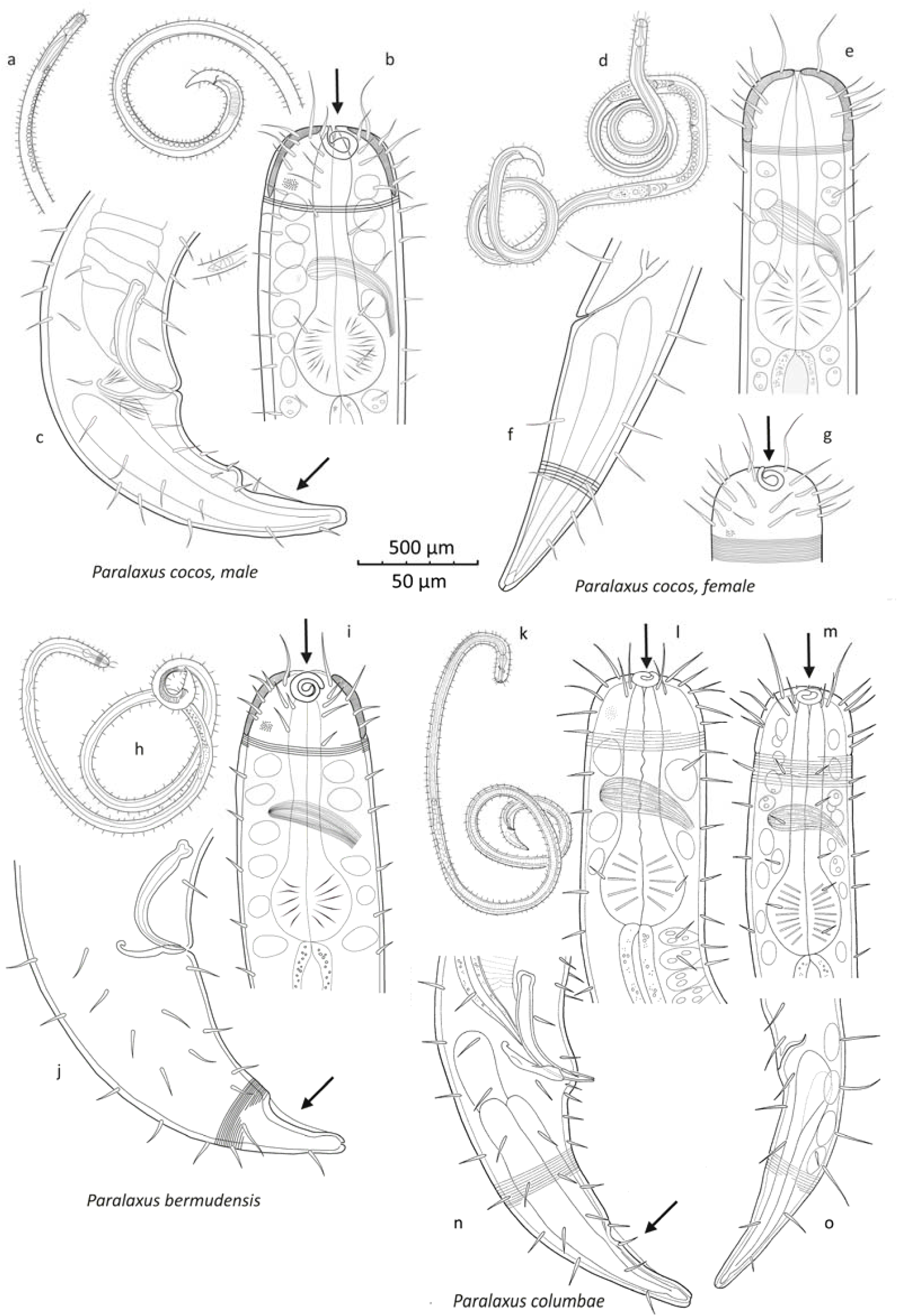
Drawings of three different *Paralaxus* species comparing their defining characters. (a - c. *Paralaxus cocos* nov. spec. male, (a) total view, (b) anterior end (c) posterior end (d – g) *P. cocos* female, (d) total view (e) anterior end, optical section (f) anterior end, surface view (h – j) *P. bermudensis* nov. spec. male (h) total view (i) anterior end (j) posterior end (k – o) *P. columbae* nov. spec. male and female. (k) total view of male paratype, (l) anterior end of male,(m) anterior end of female (n) posterior end of male (o) posterior end of female. Camera lucida drawings. Arrows point to amphidial fovea and velum on male tail.

**Fig. 3.**
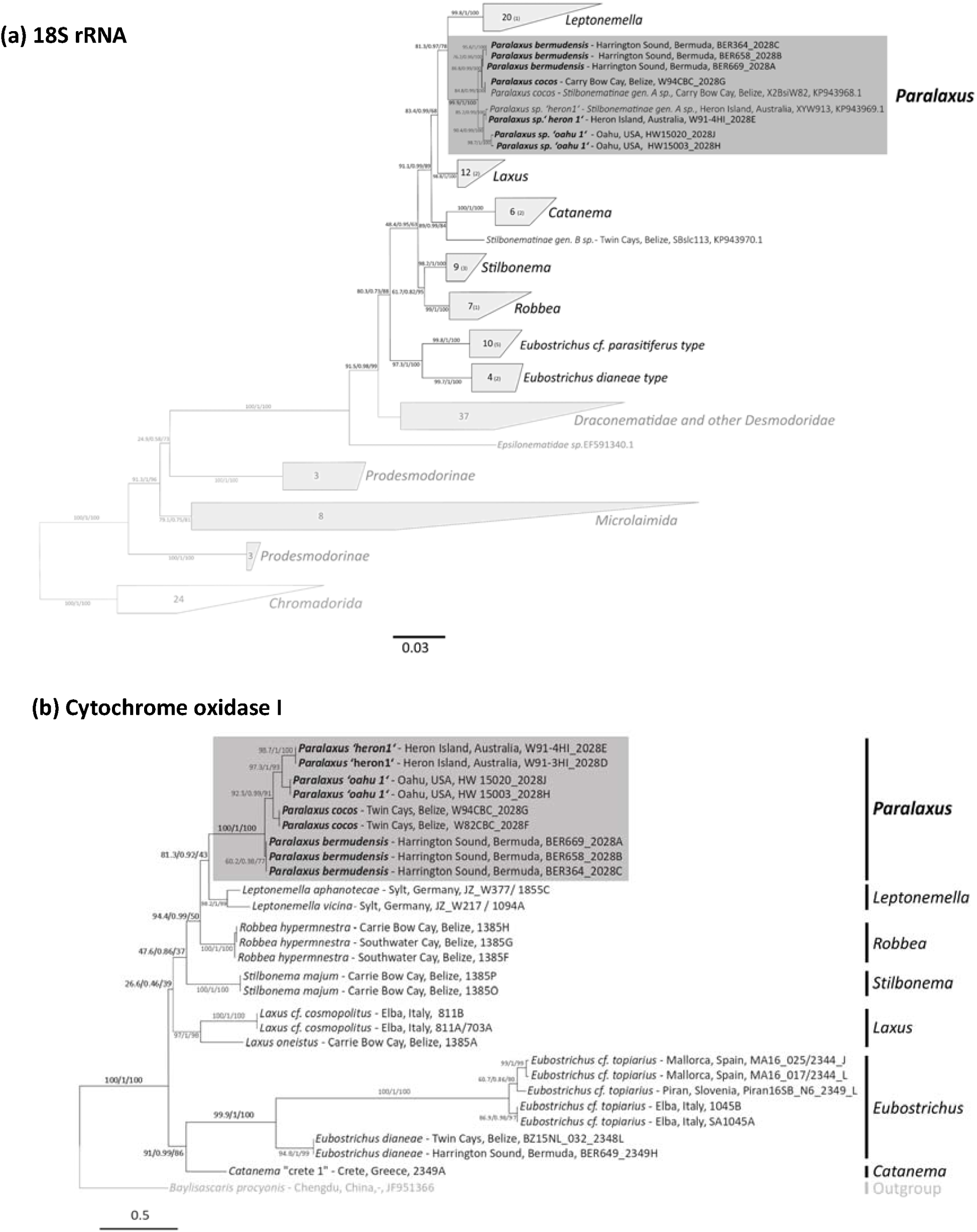
Phylogenetic relationship of Paralaxus species with other stilbonematine nematodes based on the 18S rRNA (a) and cytochrome oxidase subunit I (COI) genes (b). The trees were calculated using IQ-Tree. Support values are given in the following order: SH-aLRT support (%) / aBayes support / ultrafast bootstrap support (%). Species described in this study are highlighted in bold. Provisional working names for undescribed genera or species are given in quotes. The scale bar represents average nucleotide substitutions per site.

We describe three new species, and report on two additional ones.

#### *Paralaxus cocos* sp. nov

ZooBank registration (http://zoobank.org): urn:lsid:zoobank.org:act A659918E-FD88-4584-8115-CD4C396AE400

GenBank entries for 18S rRNA and COI sequences: see Table S1.

Species from Belize, Central America can be identified by an amphidial fovea with one and a half turns and the bulbus occupies 32 - 35% of the pharynx length. The velum of male specimens makes up 43 - 47% of the tail length and the spicula are moderately arcuate (Fig. 2a-g).

#### *Paralaxus bermudensis* sp. nov

ZooBank registration (http://zoobank.org): urn:lsid:zoobank.org:act: 8697C6A0-3508-4833-81DC-D2A86B0B00AC

GenBank entries for 18S rRNA and COI sequences: see Table S1.

Species from Bermuda can be identified by an amphidial fovea with two turns and the bulbus occupies 30 - 32% of the pharynx length. In male specimens the velum makes up 30 – 40% of the tail length and the spicula are strongly arcuate (Fig. 2h-j).

#### *Paralaxus columbae* sp. nov

ZooBank registration (http://zoobank.org): urn:lsid:zoobank.org:act:4BABABB3-1DD4-46F1-A69C-BC06C1271D01

Species from Florida Keys, USA can be identified by an amphidial fovea with 1– 1.2 turns and the bulbus occupies 29 - 33% of the pharynx length. In male specimens the velum makes up 35 - 37% of tail length and the spicula are weakly arcuate (Fig. 2k-0).

### Paralaxus species from Australia and Hawaii

GenBank entries for 18S rRNA and COI sequences see Table S1.

Representatives of two additional species, *Paralaxus* sp. “heron 1” and P. sp. “oahu 1” have been found off Heron Island, Great Barrier Reef, Australia and Oahu, Hawaii Islands. Due to the lack of properly preserved material, a formal description is currently not possible. Morphological details and measurements taken from microphotographs of live animals prior to fixation are presented in the Supplementary Information (Table S2).

Detailed *Paralaxus* species descriptions are presented in the Supplementary Information (Supplementary Results and Figs. S1 – S8).

### Key to the species

Amphidial fovea with 2 turns; bulbus 30 - 32% of pharynx length, velum 30 – 40% of tail length, spicula strongly arcuate ***Paralaxus bermudensis* sp. nov**.

Amphidial fovea with 1.5 turns, bulbus 32 - 35% of pharynx length, velum 43 - 47% of tail length, spicula moderately arcuate ***P. cocos* sp. nov**.

Amphidial fovea with 1 – 1.2 turns, bulbus 29 - 33% of pharynx length, velum 35 - 37% of tail length, spicula weakly arcuate ***P. columbae* sp. nov**.

### Morphological delineation of the genera Paralaxus, Laxus and Leptonemella

Most characters that identify the genus *Paralaxus* are shared with one or more genera of the Stilbonematinae, but their combination is unique and characteristic. *Paralaxus* is similar to the genus *Laxus* (as reflected in the genus name), but also to the genus *Leptonemella.* The three genera share the extreme forward position of the spiral amphidial fovea, a well-developed cephalic capsule, and a gubernaculum without apophysis. *Paralaxus* can be differentiated from *Laxus* by the lack of a block-layer in the cephalic capsule, the presence of a velum on the male tail and in addition by the multi-layered symbiont coat. *Paralaxus* shares the lack of the block-layer and the multilayered symbiont coat, with *Leptonemella*, but is distinguished by the greater relative pharynx length b’ (pharynx length/body diameter at end of pharynx), the presence of a velum on the male tail, and the lack of sexual dimorphism in the shape of the amphidial fovea.

A special case is *L. brevipharynx* *Armenteros et al.* (2014 b). According to phylogenetic analyses of its 18S and partial COI data (Fig. 3a and b) the species belongs to the genus *Leptonemella.* It is morphologically similar to P. cocos sp. nov., and despite being molecularly a bona-fide *Leptonemella*, has a short pharynx and no sexual dimorphism in the shape of the amphidial fovea. There are, however, two morphological features, which clearly separate *L. brevipharynx* from *Paralaxus*: the lack of a velum on the tip of the tail in males and the lack of the long forward directed cephalic setae characteristic for all *Paralaxus* species.

### Molecular data corroborates the monophyly of the subfamily Stilbonematinae and all its genera, including Paralaxus gen. nov

In addition to the nine *Paralaxus* specimens, we generated and analysed single worm metagenomes from 20 specimens covering six additional stilbonematine genera and extracted both host 18S and COI gene sequences for all specimens (Supplementary Table S1). To analyse the monophyly of the *Stilbonematinae*, we reconstructed their phylogenetic relationships based on the 18S gene, as this is the only gene with sufficient taxon sampling. The 18S gene matrix, with a minimum sequence length of 1300 bp, consisted of sequences from 78 *Stilbonematinae* as well as from 52 closely related *Desmodorida* and *Microlaimida*, and an outgroup of 24 *Chromadorida* sequences. The *Stilbonematinae* formed a highly supported clade, with *other non stilbonematine Desmodoridae* and *Draconematidae* as closest relatives (Fig. 3a).

For the analysis of the Stilbonematinae genera, we created a mitochondrial COI tree in addition to the 18S tree, using 27 metagenome-derived stilbonematine COI sequences and an ascarid as outgroup (*Baylisascaris procyonis* mitochondrion, JF951366) (Fig. 3b). Based on both datasets, all Stilbonematinae genera including *Paralaxus* gen. nov. formed well-supported clades (Fig. 3a and b), thus corroborating our morphological assignments. In both the 18S and the COI-based analysis, the genus *Leptonemella* was phylogenetically most closely related to *Paralaxus*, with high statistical support in the 18S dataset and acceptable support in the COI-based phylogenies (Fig. 3a and b). Other than that, the branching patterns between genera were largely inconsistent between the COI and 18S datasets. This could be linked to the remarkably low support values for many internal nodes in the COI dataset compared to the much higher spectrum of support values for the 18S-based tree topologies.

In an extended 18S and COI data-matrix, we also included eight short-length, PCR-amplified sequences of five different *Stilbonematinae* species with a record of problematic phylogenetic placement – *Laxus parvum, Robbea porosum, Stilbonema brevicolle, Catanema exile* and *Leptonemella brevipharynx* (Armenteros et al. 2014b) (Supplementary Figs. S9 and S10). With our new 18S and COI-based datasets the placements of *Leptonemella brevipharynx* and *Catanema exile* sequences were resolved and both clustered with sequences of their respective genera. For the sequences of the other three species we observed inconsistent grouping with *Stilbonematinae* genera between the datasets (Supplementary Figs S9 and S10). In the extended 18S tree, two *Stilbonema brevicolle* sequences appeared in two different clades, one closely related to *Leptonemella*, while the other one clustered with a yet undescribed genus-level clade designated as genus ‘B’ (Zimmermann et al. 2016). In contrast to the 18S-based placement, the *Stilbonema brevicolle* sequences grouped together in the extended COI tree and formed a novel sister clade to several genera including *Stilbonema, Robbea, Leptonemella* and *Paralaxus* (Supplementary Fig. S10). In the extended 18S data-matrix, the *Laxus parvum* sequences clustered within the genus *Robbea* (Supplementary Fig. S9). Similarly, in the COI dataset the *L. parvum* sequence grouped with the only available sequence of the genus *Robbea* (Supplementary Fig. S10). The sequences designated as *Robbea porosum* formed a divergent genus-level clade but the placement of this clade was not consistent for the two marker genes (Supplementary Figs S9 and S10). In the 18S-based analyses they were the sister clade to the genus *Laxus*, while in the COI dataset they formed the sister clade to all other Stilbonematinae.

### Symbiont 16S phylogeny shows genus-level specificity

We constructed a matrix from all full-length 16S ‘Thiosymbion’ sequences, available in the databases and 21 metagenome-derived symbiont sequences - using five free-living Chromatiaceae as an outgroup. All symbiont sequences from the same host genus clustered in well-supported clades, irrespective of the number of gutless oligochaete symbionts included within those clades (Fig. 4). In congruence to the host data, the symbionts of all newly sequenced *Paralaxus* individuals as well as the two previously published sequences from the Caribbean and Australia (Zimmermann et al. 2016) formed a supported clade in the 16S phylogeny (Fig. 4).

**Fig. 4.**
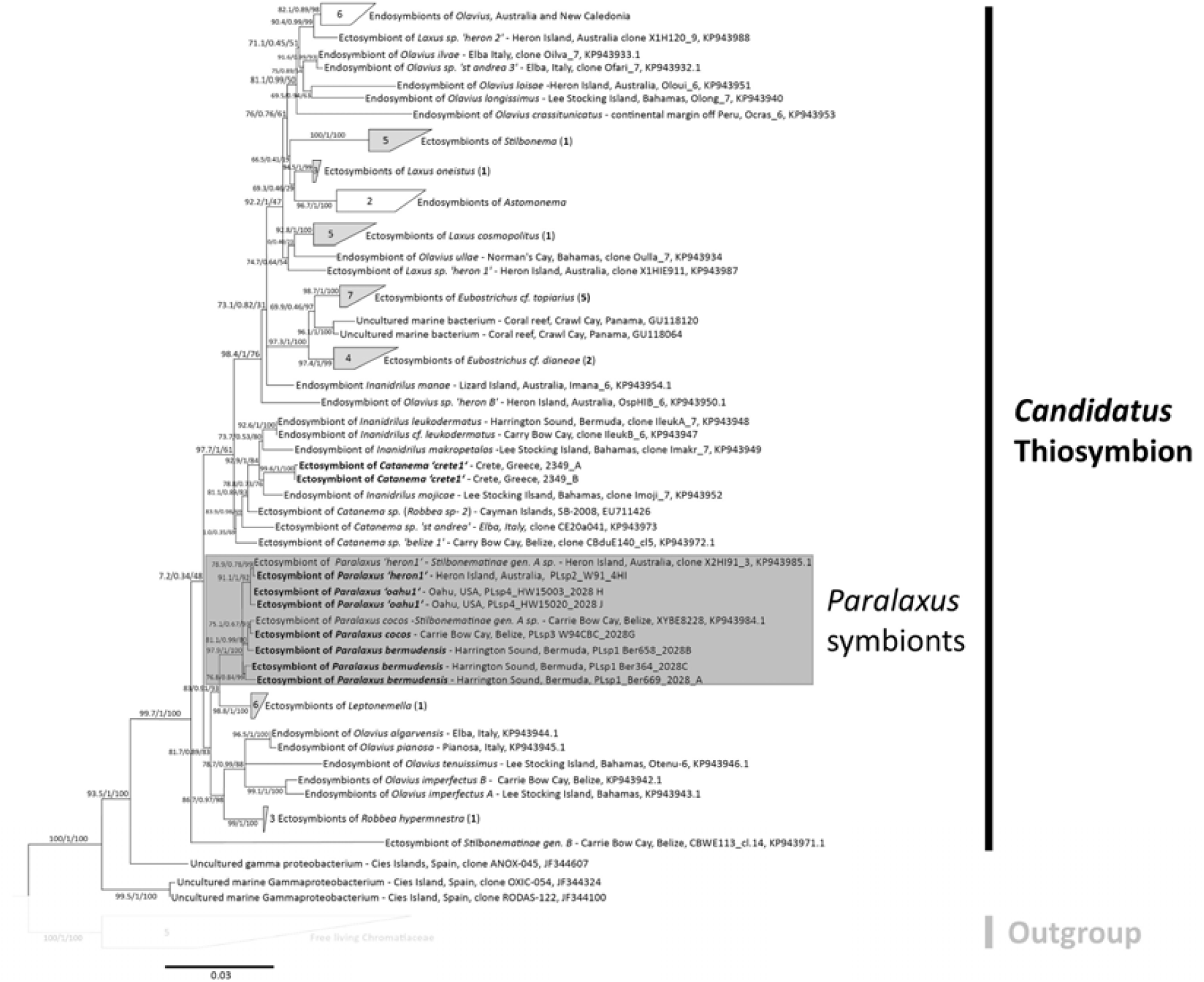
*Candidatus* Thiosymbion phylogeny, including symbionts of *Paralaxus* species and other stilbonematine nematodes, based on the 16S rRNA gene. The tree was calculated using IQTree and node support is given in the following order: SH–aLRT support (%) / aBayes support / ultrafast bootstrap support (%). Symbionts of species described in this study are marked bold or highlighted in bold numbers next to collapsed branches. Provisional working names for undescribed genera or species are given in quotes. The scale bar represents average nucleotide substitutions per site.

### COI primers cannot cover the compositional diversity in Stilbonematinae

Despite a highly sensitive PCR setup, we could not amplify the COI of any *Paralaxus* specimen using the widely used primer set by Folmer et al. (1994) (data not shown). To elucidate these PCR-based amplification problems, we analysed all our metagenome-derived full-length stilbonematine nematode COI sequences for compositional patterns. The COI sequences of the 19 stilbonematine nematode species that belonged to seven genera had a highly variable guanine and cytosine (GC) content, ranging from 27.7% in the genus Robbea to 53.1 % in the genus *Leptonemella.* We identified substantial priming problems with the Folmer et al. primer set as well with as a second set (JB3F and JB5R) that is commonly used for COI amplification (Bowles et al. 1992, Derycke et al. 2010, Armenteros et al. 2014a,b). Across the *Stilbonematinae* diversity the Folmer et al. primers had a range of 3 - 11 mismatches for the 1490F primer and 1 - 5 mismatches for the 2198R primer. Similarly, the primers JB3F and JB5R used in a recent study by Armenteros et al. (2014b) showed up to 11 mismatches for each primer (Fig. 5). Of all four primers tested only the JB5R perfectly matched, and only to three sequences from Robbea *hypermnestra* (Fig. 5). We performed an *in-silico* analysis that mimics PCR conditions of low stringency, which would overcome up to 11 primer mismatches and would cover the whole diversity of *Stilbonematinae.* Our analysis showed that such permissive conditions would lead to unspecific binding at multiple alternate sites of the COI gene. We tried to design alternative primer sets, but no primer pair, neither for the full-length nor for partial regions, would reliably target the COI gene and would cover all genera of *Stilbonematinae.*

**Fig. 5.**
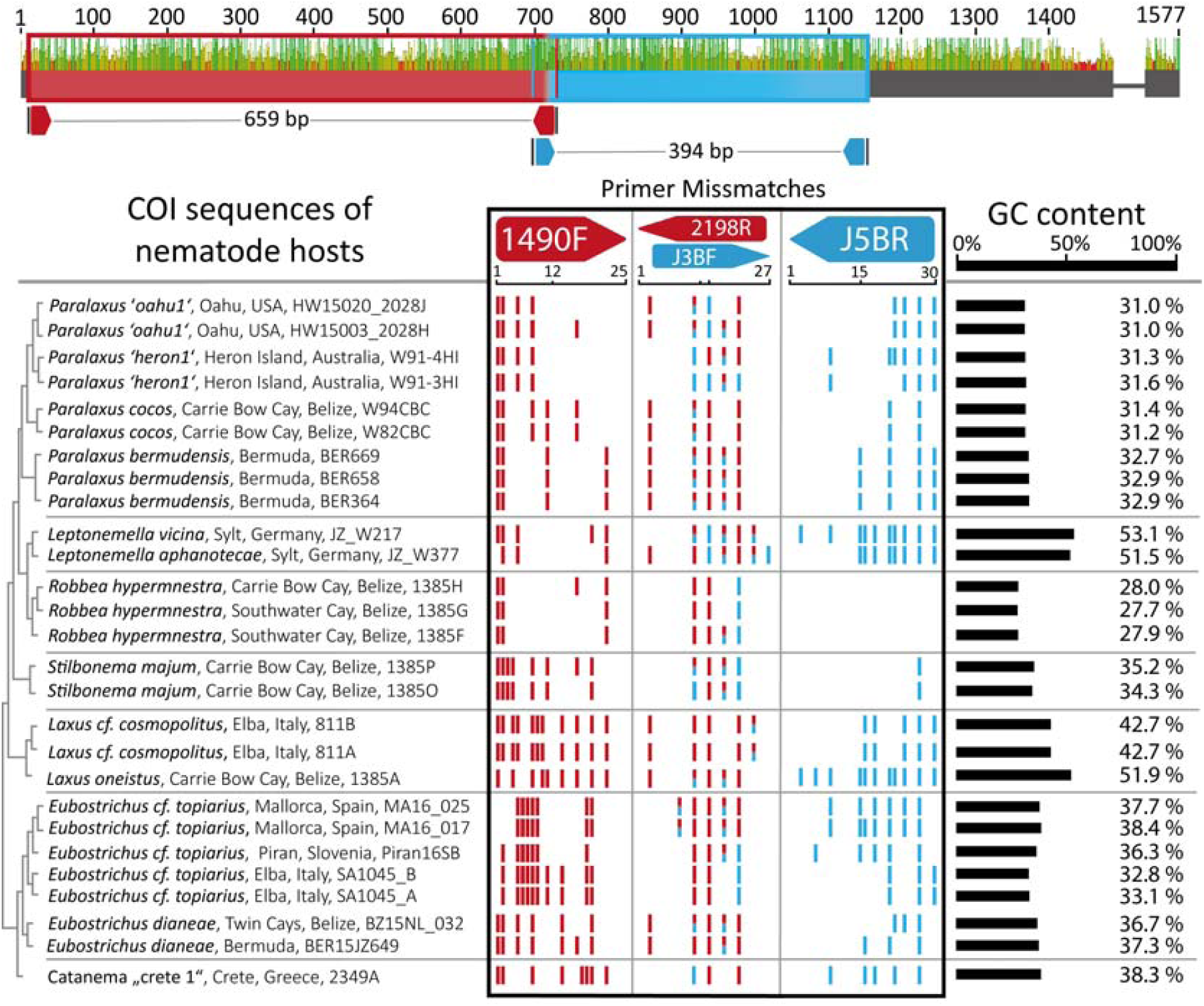
Primer mismatch analysis of the two commonly used COI primer sets. Potential priming sites (forward and reverse) for the primer sets 1490F/ 2198R (Folmer *et al.* 1994) and J3BF/ J5BR (Armenteros et al. 2014a,b) along the COI gene are shown in red and blue on the top. The number of mismatches for each primer, when matched to metagenome– extracted full–length COI sequences from seven representative stilbonematine nematode genera, is shown below on the left. Average GC content for each full–length COI sequence is represented by the scale bar on the right.

### The symbiont coat as a new diagnostic feature for Stilbonematinae genera

In addition to several host-based morphological differences, the presence of a multi-layered coat distinguishes the new genus *Paralaxus* from *Laxus*, which carries a monolayer of symbionts. Extending from this observation we analysed the structure and morphology of the bacterial coat of all *Stilbonematinae* genera and found substantial and highly informative differences. We therefore propose to include the description of the bacterial coat given below into the genus trait sets as a new character (see also iconographic overview of the different genera in Fig. 6).

**Fig. 6.**
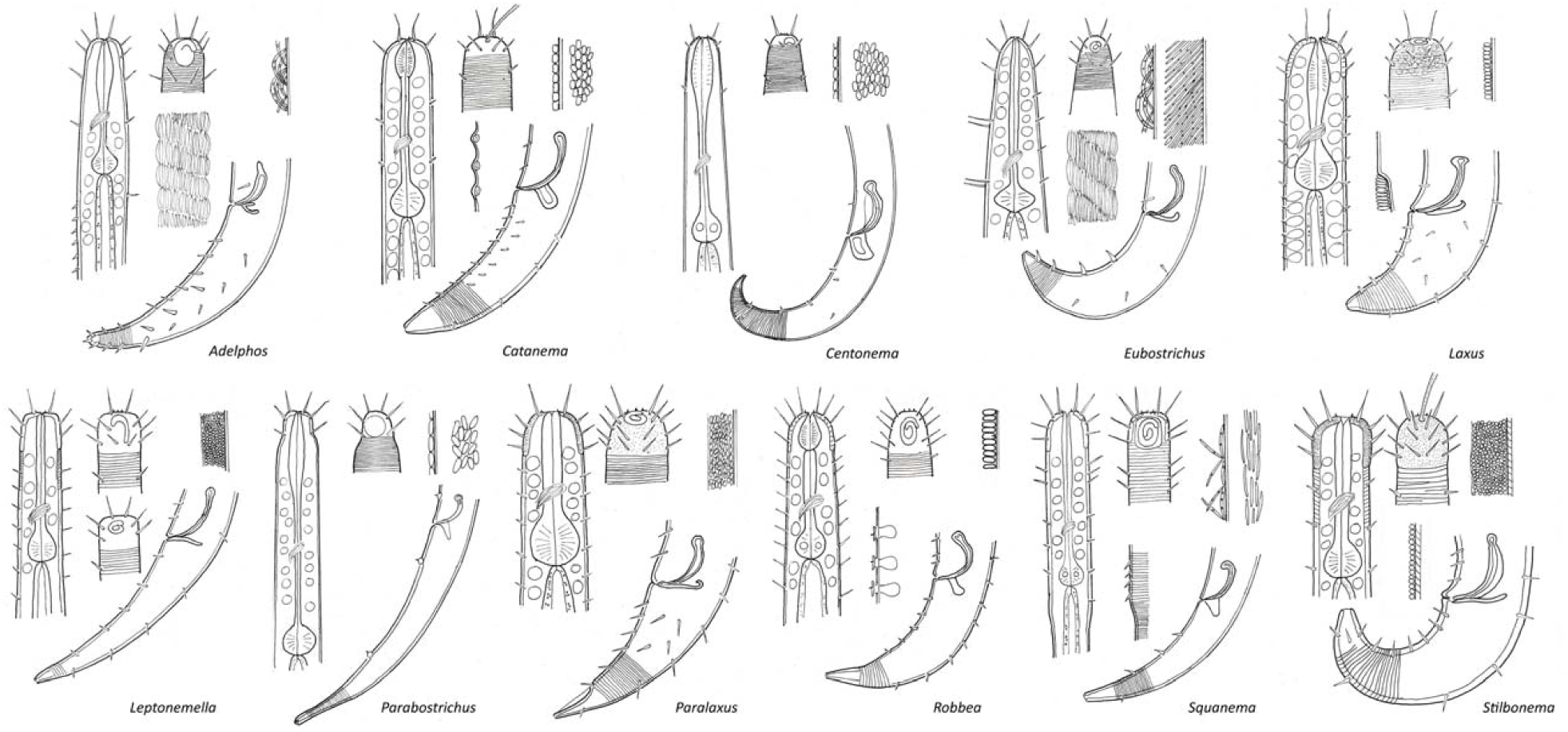
Cellular morphology and coat structure of symbiotic bacteria on different stilbonematine nematode genera. All described genera of Stilbonematinae including *Paralaxus* nov. gen. are depicted in alphabetical order. Each pictogram shows the anterior end of a male in optical section (left), the head region of a male in surface view (top center), male posterior region with spicular apparatus (lower right) and shape and arrangement of symbionts (upper right). Special features such as arrangement of complex symbiont coat (*Adelphos, Eubostrichus*), male supplementary structures (*Catanema, Robbea, Stilbonema*), sexual dimorphism in the shape of the amphidial fovea (*Leptonemella*) or change in cuticular structure (*Laxus, Squanema*) are shown in the center of the pictogram. Drawings do not represent a defined species but include the general characters of each genus.

> *Adelphos* Ott 1997
>
> Crescent-shaped bacteria, all cells in contact with host cuticle, double-attached, forming complex monolayer, covering whole body except anterior and posterior tip
>
> *Catanema* Cobb 1920
>
> Cocci or coccobacilli, all cells in contact with host cuticle, forming simple monolayer, covering whole body except anterior and posterior tip or starting at distance from anterior end
>
> *Centonema* Leduc 2013
>
> Rods (bacilli), lying parallel to cuticle, all cells in contact with host cuticle, forming simple monolayer, probably covering whole body
>
> *Eubostrichus* Greef 1869
>
> *E. parasitiferus-Type*
>
> Crescent-shaped bacteria, all cells in contact with host cuticle, double-attached, forming complex monolayer, covering whole body except anterior and posterior tip
>
> *E. dianeae-Type*
>
> Filamentous bacteria, all cells in contact with host cuticle, singly attached, forming complex monolayer, covering whole body except anterior and posterior tip
>
> *Laxus* Cobb 1894
>
> Cocci, coccobacilli or rods (bacilli), all cells in contact with host cuticle, rods standing perpendicular to cuticle surface, forming simple monolayer, covering whole body except anterior and posterior tip or starting at distance from anterior end
>
> *Leptonemella* Cobb 1920
>
> Cocci, only a fraction of cells in contact with host cuticle, forming multilayer embedded in gelatinous matrix, covering whole body except anterior and posterior tip
>
> *Parabostrichus* Tchesunov, Ingels & Popova 2012
>
> Crescent shaped bacteria, all cells in contact with host cuticle, double-attached, forming complex monolayer, covering whole body except anterior and posterior tip
>
> *Paralaxus*
>
> Cocci or coccobacilli, only a fraction of cells in contact with host cuticle, forming multilayer embedded in gelatinous matrix, covering whole body except anterior and posterior tip
>
> *Robbea* Gerlach 1956
>
> Cocci, coccobacilli or rods (bacilli), all cells in contact with host cuticle, rods standing perpendicular to cuticle surface, forming simple monolayer, covering whole body except anterior and posterior tip or starting at distance from anterior end
>
> *Squanema* Gerlach 1963
>
> Crescent-shaped bacteria, all cells in contact with host cuticle, single or double attached, forming monolayer, starting at distance from anterior end
>
> *Stilbonema* Cobb 1920
>
> *S. brevicolle-Type*
>
> Cocci or rods, all cells in contact with host cuticle, forming simple monolayer covering whole body except for anterior and posterior tip
>
> *S. majum-Type*
>
> Cocci, only a fraction of cells in contact with host cuticle, forming multilayer embedded in gelatinous matrix, covering whole body except anterior and posterior tip

## Discussion

### COI-based barcoding is not suitable for Stilbonematinae

The COI gene is the most common marker gene for PCR-based animal (meta) barcoding, despite the observation that highly diverse base composition have led to unreliable amplification results in many animal phyla including nematodes (Bhadury et al. 2006, Derycke et al. 2010, Deagle et al. 2014). By utilizing single nematode metagenomics we could overcome such amplification problems and show that the COI is a valuable phylogenetic marker gene to distinguish stilbonematine genera and species. However, analyses of the metagenome-derived full-length COI sequences revealed that within *Stilbonematinae* the COI gene has a highly diverse base composition, prohibiting the design of specific primers on any region of the COI gene. Our mismatch analyses for commonly used primer sets showed multiple mismatches as well as possible alternate binding sites. One may overcome priming mismatches by using very low stringency PCR conditions, at the cost of specificity. Armenteros et al. (2014b) were able to amplify COI fragments from stilbonematine DNA extracts but morphological species descriptions and 18S-based results did not conform to the phylogenetic placement based on the COI. One possible explanation could be that the PCR picked up minimal contaminations of other *Stilbonematinae* with lower mismatch counts, instead of the gene of the target organism with high numbers of mismatches (Fig. 5). Together these factors show that the COI gene is not well suited for PCR-based barcoding in *Stilbonematinae.*

### Metagenomics-based molecular data leads to robust phylogenies of Stilbonematinae

Up to this study, 18S and COI genes for *Stilbonematinae* were PCR-amplified from DNA extracts – a technique known to be highly susceptible to contamination and bias. This is reflected in two studies by Armenteros et al. (2014a,b), which contain phylogenies based on short PCR-amplified stilbonematine nematode sequences. These studies casted doubt on the monophyly of the subfamily *Stilbonematinae*, as well as the validity of several stilbonematine genera. As discussed by the authors themselves (Armenteros et al. 2014a) and Leduc and Zhao (2016), these conflicting results could have been caused by misidentification of species, PCR bias or artefacts of the phylogenetic reconstruction. To resolve these questions, we used single worm metagenomics from photo-documented specimens to generate high quality and full-length 18S and COI sequences that cover the majority of the described stilbonematine genera. Our dataset corroborated the monophyly of the subfamily *Stilbonematinae* and verified the taxonomic status of all analysed stilbonematine genera. We caution against the use of short and PCR-based stilbonematine nematode 18S and COI sequences (see Supplementary Figs. S9 and S10) for phylogenetic analyses and barcoding.

### Low coverage next generation sequencing should replace PCR-based barcoding approaches

Instead of using PCR-based barcoding, a straightforward solution to capture marker genes of a target organism are single individual genome or metagenome sequencing approaches employing next generation sequencing (NGS). Our data shows that an NGS-based approach has the advantage that not only single genes but a wide range of marker genes, such as the full rRNA operon as well as the full mitochondrial genome can be recovered from shallow sequencing depths. Recent studies have shown that successful library preparation can be done from as little as 1 picograms of DNA (Rinke et al. 2016) and has been used on a range of microscopic eukaryotic taxa (Jäckle et al. 2018, Gruber-Vodicka et al. 2019, Seah et al. 2019). Commercially available kits allow input amounts as low as 1 ng of DNA, and have been used for low cost library construction protocols (Therkildsen & Palumbi 2017). Low coverage NGS-based genomic sequencing could even allow for population genetic studies when combined with appropriate replication and probabilistic allele calling techniques (Therkildsen & Palumbi 2017). Additionally, the associated microbiome is also sequenced, which opens many new avenues for host-microbe research.

### The symbiont coat is an additional character of host taxonomy

In microscopic organisms such as marine meiofauna, taxonomic characters are often inconspicuous and not easily recognized. With their relatively simple body plan only few morphological features are the defining characters for nematode genera. For *Stilbonematinae* this leads to the problematic situation that characters overlap and only combinations of multiple characters can delineate genera. This was especially evident in the case of *Paralaxus* gen. nov., as outlined above. The most conspicuous feature of *Stilbonematinae* is their coat, but initial descriptions of both genera and species often interpreted the obligate symbiotic coat as part of the host (Greeff, 1867), as parasites (Chitwood, 1936) or disregarded the symbiotic coat completely (Cobb, 1920). In his taxonomic review Tchesunov (2013) gives brief descriptions of their cellular morphologies but does not mention the more informative arrangement of the coat itself.

The fact that *Paralaxus*, but also all other stilbonematine host genera are associated with a specific clade of symbiotic bacteria (based on 16S sequences) (Fig. 4 and Zimmermann et al. 2016) allows the use of the symbionts’ cellular morphology and coat structure as taxonomic characters for the holobiont. Both cell morphology and coat structure proved to be characteristic for host taxa at the genus level (Fig. 6). Interestingly, in the stilbonematine genus *Eubostrichus*, species can host symbionts of either crescent or filamentous shapes (Fig. 6). The 18S and COI sequences of hosts harbouring the two morphotypes are phylogenetically distinct (Fig. 3a and b), and this clustering is also reflected in the symbiont phylogeny (Fig. 4). This suggests that the genus *Eubostrichus* could require a taxonomical reinvestigation. A similar case can be made for the genus *Stilbonema* were at least two coat arrangements (Fig.6) are present but comprehensive molecular data does not exist yet. These two examples emphasize the taxonomic value of the symbiotic coat to reassess the assignment of specimens to genera. Furthermore, coat-based characters are easily recognized in live specimens as well as in formaldehyde-preserved material where the coat is often intact. The coat thus facilitates stilbonematine genus level differentiation during stereomicroscopic investigations, e.g. directly in the field. We emphasize that the use of the symbiont coats as additional character will both simplify and enhance *Stilbonematinae* identification and taxonomy.

## Conclusions

The last decade has revealed the ubiquitous and intimate link between animals and the microbial world. Conspicuous holobiont features such as the stilbonematine nematode symbiont coats not only support taxonomic identification, but also emphasize the co-evolution of the symbioses. The highly variable base composition of the *Stilbonematinae* mitochondrial COI hints at the possible coupling between the obligate and long-term symbiont and host mitochondrial genomes. These findings also reiterate the importance to analyse more than a single marker gene. We are entering a new age in invertebrate zoology – it is becoming a viable option to employ data-intensive NGS sequencing for multi-locus analyses and, when performed in a shotgun metagenomic approach, this allows to consider the permanently associated microbiome at the same time. Exploring these options, our results show how metagenomics and the integration of holobiont features can shape taxonomic studies and likely many more fields of zoology in the future.

## Acknowledgements

We are grateful to the excellent assistance of the staff and for laboratory facilities provided by the Carrie Bow Cay Marine Laboratory (Smithsonian Institution) in Belize, the Heron Island Research Station in Australia, the Bermuda Institute of Ocean Sciences in St. George’s, the HYDRA Institute on Elba, Italy and the Wadden Sea Station Sylt of the Alfred Wegener Institute in Bremerhaven, Germany. We thank Ramon Rosello-Mora and the IMEDEA staff for field work support on Mallorca, Spain. We further gratefully acknowledge the collaboration of the Bermuda Aquarium in Flatts Village for granting us access to the sampling site and their facilities. We thank Nicole Dubilier for funding HGV, JZ and NL through the Max Planck Society and the UNI:DOCs program at the University of Vienna for funding FS.

We thank CIUS (Core Facility Imaging and Ultrastructural Research) at the Faculty of Life Sciences, University of Vienna, Austria and especially Daniela Gruber for assistance with electron microscopy. Many thanks to Silke Wetzel, Manuel Kleiner and Cécilia Wentrup for their contributions during the Bermuda and Belize sampling expeditions. This is contribution XXX from the Carrie Bow Cay Laboratory (CCRE program, Smithsonian Institution). Completion of the final version of the manuscript was supported by grant P 31594 (J. Ott, PI) of the Austrian Science Fund (FWF).

## Supplementary Information

### SI Results

#### Descriptions

Class Chromadorea Inglis, 1983

Subclass Chromadoria Pearse, 1942

Order Desmodorida De Coninck, 1965

Suborder Desmodorina De Coninck, 1965

Superfamily Desmodoroidea Filipjev, 1922

Family Desmodoridae Filipjev, 1922

Subfamily Stilbonematinae Chitwood, 1936

#### *Paralaxus* gen. nov

http://zoobank.org: 417691DB–5EE3–49B7–B838–1BBF1569235F

#### Diagnosis

Stilbonematinae. Cuticle transversely striated, except for the head region and the tip of the tail. The striation is coarser in the anterior region, becoming finer at 3 – 4 pharynx lengths. Cephalic capsule distinct, without a block–layer, finely punctated; cephalic setae as long or longer than the sub–cephalic setae, usually directed straight forward. Amphidial fovea an open spiral, ventrally wound, in extreme forward position. No sexual dimorphism in the shape of the amphidial fovea. Pharynx bulbus large, muscular. Gubernaculum dorsally directed with terminal hook, without apophysis. Males have a distinct velum at the tip of the tail. Glandular sense organs (gso) enlarged ventrally in post–pharyngeal region in males; anterior and posterior to vulva in females. Body covered by a multi–layered coat of symbiotic coccobacilli.

Etymology: Superficially resembling species of the genus *Laxus* Cobb.

Type species: *Paralaxus cocos* sp. nov.

#### Symbiont Diagnosis

The bacterial coat consists of coccobacilli shaped cells between 0.8 µm to 1.7 µm in size, which are distributed over the whole body, leaving only the tip of the tail and the cephalic capsule uncovered. They are arranged in a thick multilayer from 6 up to 12 layers of bacteria, which are embedded in a gelatinous matrix showing no layer separation or structure. Only bacterial cells at the bottom of the coat are in contact with the cuticle of the nematode host. Most cells show pili–like structures that are connected to the worm surface and/or to neighboring bacteria. The bacterial cells divide transversely.

#### *Paralaxus cocos* sp. nov

(Figs. 2 – 23)

ZooBank registration: http://zoobank.org A659918E–FD88–4584–8115–CD4C396AE400

#### Type material

Holotype (male), 3 paratypes (male), 3 paratypes (female), 4 paratypes (juvenile), deposited at the Natural History Museum Vienna (accession numbers NHMWZooEvMikro 5689 – 5699)

#### Measurements

See Table 1

#### Additional material

Several specimens in the collection of JAO and those used for SEM.

#### Type locality

Subtidal fine sand among Red Mangrove (*Rhizophora mangle*) roots and turtle grass (*Thalassia testudinum*), 10 – 50 cm depth, W–side of Twin Cayes, BELIZE (16°49’45.6”N 88°06’28.9”W)

#### Distribution

Regularly in fine to medium subtidal sands around Carrie Bow Cay, Curlew Cay and Twin Cayes (Belize).

#### Etymology

From the Latin name of the Coconut Tree, *Cocos nucifera*, one of which played an important role in the research trip that led to the discovery of the new genus.

### Description

Body cylindrical, head diameter at level of cephalic setae 20 – 25 µm, diameter at level of posterior margin of amphidial fovea 23 –28 µm in males, 20 – 25 µm in females, at end of pharynx 46 – 57 µm, maximum body diameter 46 – 66 µm, anal diameter 50 – 55 µm in males, 40 – 42 µm in females. Tail conical, 115 – 120 µm long in males and 98 – 125 µm in females.

Cuticle transversely striated except for the first 25 – 30 µm of the head and the last 40 – 50 µm of the tail (Fig. 14), striae 0.6 to 0.7 µm wide (14 – 16 striae/10 µm) in anterior body part, 0.4 –0.5 µm (20 – 23 striae/10µm) after 3 – 3.7 pharynx length from anterior end. The anterior most circle of head sensillae (inner labial sensillae) is represented by 6 spoon–shaped papillae within the mouth opening, in lateral, subventral and subdorsal position. The second circle consists of 6 short outer labial sensillae, 1 µm long, on the margin of the membranous buccal field in lateral, subventral and subdorsal position; 4 cephalic setae near the anterior margin of the amphidial fovea, 20 – 28 µm long; 3 circles of 8 subcephalic setae each, 15 – 20 µm long, on the non– striated part of the head region, the submedian rows with additional short setae, 10 µm long; 8 rows of somatic setae along the whole length of the body, 12 µm long, spaced 20 µm apart. Non–striated part of the tail in females with 2 pairs of setae, in males with a velum occupying 43 – 47% of tail length. Three caudal glands extending to the gubernaculum. Amphidial foveas situated at the anterior end bordering the buccal field, an open spiral, ventrally wound, with 1.5 turns, 10 – 15 µm long and 14 – 15µm wide in males, smaller in females, 6 – 8 µm long and 11 – 12 µm wide. In one male specimen the amphidial fovea was dorsally wound (Fig. 3).

Pharynx 89 – 130 µm long, with a minute buccal cavity, 5 µm long, leading into a slightly dilated corpus, which is only indistinctly separated from the isthmus. Posterior bulbus spherical, muscular, large, 28 – 43 µm in diameter. No cardia. The bulbus to pharynx ratio is 32 – 35% in males and 30 – 34% in females.

Nerve ring 45 – 55 µm from anterior end; no secretory–excretory pore or ventral gland seen; 8 rows of glandular sense organs (two in each lateral and each median line).

Males monorchic, testis on the right side of the intestine, beginning at about 36–37% of body length; spicula strong, arcuate, cephalate proximally, 50 – 51 µm (chord) or 60 – 65 µm (arc) long; gubernaculum straight, corpus embracing spicules, 31 – 35 µm long, end hook–shaped. Sperm globular. Row of enlarged gso along the mid–ventral line 400 µm long, beginning at 250 µm from the anterior end.

Females didelphic, ovaries reflexed; vulva at 44 – 45% of body length. Enlarged ventral gso anterior and posterior to vulva, extending 250 µm in either direction.

#### *Paralaxus bermudensis* sp. nov

(Figs. 24 – 30)

ZooBank registration: http://zoobank.org 8697C6A0–3508–4833–81DC–D2A86B0B00AC

#### Type material

Holotype (male), 1 paratype (male), 2 paratypes (juvenile), deposited at the Natural History Museum Vienna (accession numbers NHMWZooEvMikro 5700 – 5703)

#### Measurements

See Table 1

#### Type locality

Subtidal sand, 3 m depth, NW of Nonsuch Island, BERMUDA (32°20’ 54’’ N, 64° 39’ 54’’ W)

#### Distribution

Also found in subtidal sand in Harrington Sound and Baylis Bay, Bermuda.

#### Etymology

From the Bermuda Islands

### Description

Body cylindrical, head diameter at level of cephalic setae 25 – 35 µm, diameter at level of posterior margin of amphidial fovea 45 – 50 µm, at end of pharynx 60 – 65 µm, maximum body diameter 60 – 65 µm, anal diameter 60 – 65 µm. Tail conical, 105 – 125 µm long.

Cuticle transversely striated except for the first 30 – 35 µm of the head and the last 40 – 45 µm of the tail, striae 1 – 1.1 µm wide (9 – 10 striae/ 10 µm) in anterior body part, appr. 0.5 µm (20/ 10 µm) after 3.2 – 3.4 pharynx lengths from anterior end. The anterior most circle of head sensillae (inner labial sensillae) not seen, the second circle as in *P. cocos*; 4 cephalic setae near the anterior margin of the amphidial fovea, 20 – 25 µm long; 3 circles of 8 subcephalic setae each, 12 – 20 µm long, with additional short setae, 10 µm long on the non–striated part of the head region; 8 rows of somatic setae along the whole length of the body, 8 – 10 µm long spaced 20 µm apart. Tip of tail in males with a velum occupying 36 – 40% of tail length. Three caudal glands extending to the gubernaculums. Amphidial foveas with 2 turns, 10 – 12 µm long and 12 – 15 µm wide.

Pharynx 116 – 125 µm long, with a minute buccal cavity, corpus slightly dilated. Bulbus spherical, muscular, large, 35 – 40 µm in diameter, no cardia. The bulbus to pharynx ratio is 31 – 33%.

Nerve ring 55 – 65 µm from anterior end; no secretory–excretory pore or ventral gland seen; 8 rows of glandular sense organs (two in each lateral and each median line). Males monorchic, testis on the right side of the intestine, beginning at about 36–37% of body length; spicula strongly arcuate, cephalate proximally, 43 – 45 µm (chord) or 57 – 63 µm (arc) long; gubernaculum straight, 30 – 33 µm long, end hook–shaped. Row of enlarged GSO beginning at end of pharynx and extending 250 µm backwards. No female found.

#### *Paralaxus columbae* sp. nov

(Figs. 31 – 39)

ZooBank registration: http://zoobank.org 4BABABB3–1DD4–46F1–A69C–BC06C1271D01

#### Type material

Holotype (male), 1 paratype (male), 3 paratypes (female), 2 paratypes (juvenile), deposited at the Natural History Museum Vienna (accession numbers NHMWZooEvMikro 5704 – 5710)

#### Measurements

See Table 1

#### Additional material

Several specimens in the collection of JAO.

#### Type locality

Subtidal sand, 2 m depth, N–side of Pigeon Key, Florida, USA (24° 42’ 19’’ N, 81° 09’ 20’’ W)

#### Distribution

Only known from type locality.

#### Etymology

From the Latin *columba* (pigeon), referring to the type locality.

### Description

Body cylindrical, head diameter at level of cephalic setae 32 – 40, diameter at level of posterior margin of amphidial fovea 40 – 50 µm, at end of pharynx 54 – 55 µm in males, 55 – 60 µm in females, maximum body diameter 55 µm in males, 55 – 60 µm in females, anal diameter 50 – 55 µm. Tail conical, 108 – 115 µm long in males and 100 – 120 µm in females.

Cuticle transversely striated except for the first 30 – 35 µm of the head and the last 40 – 50 µm of the tail, striae 0.6 to 0.7 µm wide (14 – 16 striae/10 µm) in anterior body part, 0.4 –0.5 µm (20 – 23 striae/10µm) after 3 – 3.2 pharynx length from anterior end. The anterior most circle of head sensillae (inner labial sensillae) not seen. The second circle as in *P. cocos*; 4 cephalic setae at the level of the amphidial fovea, 25 – 30 µm long; 3 circles of 8 subcephalic setae each, 15 – 20 µm long, on the non–striated part of the head region, the submedian rows with additional short setae, 10 µm long; 8 rows of somatic setae along the whole length of the body, 7 – 10 µm long, spaced 15 – 20 µm apart. Males with a velum occupying 35 – 37% of tail length. Three caudal glands extending to the gubernaculum. Amphidial foveas situated at the anterior end, an open spiral with 1 – 1.2 turns, ventrally wound, 12 – 13 µm long and wide in both sexes.

Pharynx 100 – 110 µm long, buccal cavity minute, 5 µm long, corpus not dilated. Posterior bulbus subspherical, muscular, large, 30 – 35 µm long and up to 45 µm wide. No cardia. The bulbus to pharynx ratio is 29 – 33% in both sexes.

Nerve ring 45 – 55 µm from anterior end; no secretory–excretory pore or ventral gland seen; 8 rows of glandular sense organs (two in each lateral and each median line). Males monorchic, testis on the right side of the intestine, beginning at about 36–37% of the body length; spicula weakly arcuate, cephalate proximally, 50 – 54 µm (chord) or 57 – 60 µm (arc) long; gubernaculum straight, corpus embracing spicules, 31 – 33 µm long, end hook–shaped. Enlarged GSO along the mid–ventral line beginning at end of the pharynx and extending 300 – 350 µm backwards.

Females didelphic, ovaries reflexed; vulva at 44 – 48% of body length. Enlarged ventral gso anterior and posterior to vulva extending approximately 200 µm in either direction.

### Key to the species

Amphidial fovea with 2 turns; bulbus 30 – 32% of pharynx length, velum 30 – 40% of tail length, spicula strongly arcuate

#### *Paralaxus bermudensis* sp. nov

Amphidial fovea with 1.5 turns, bulbus 32 – 35% of pharynx length, velum 43 – 47% of tail length, spicula moderately arcuate

#### *P. cocos* sp. nov

Amphidial fovea with 1 – 1.2 turns, bulbus 29 – 33% of pharynx length, velum 35 – 37% of tail length, spicula weakly arcuate

#### *P. columbae* sp. nov

#### *Paralaxus* species from Australia and Hawaii

Two additional *Paralaxus* species, one from Australia and one from Hawaii have been discovered by sequencing their 18S rRNA and COI genes alone. Due to the lack of properly preserved material, a formal description is therefore currently not possible. However, we were able to make measurements and morphological descriptions from microphotographs that were taken from the live animals prior to sequencing, which are listed below:

#### *Paralaxus* sp. “heron 1””

(Figs. 40 – 42)

#### Sample location

Heron Island, Great Barrier Reef, AUSTRALIA (24° 26’ 36’’ S, 151°54’ 47’’ W)

#### Material

1 female

#### Description

(measurements taken from LM micrographs of sequenced specimen) Since the total length could not be obtained from the micrographs, the parameters a, b and c could not be calculated. Maximum diameter at the level of a fully developed egg was 86 µm, in other regions of the body 78 µm, which was also the diameter at the end of the pharynx. Cuticle striated, striae 1.2 to 1.3 µm wide. Pharynx 122 µm long, bulbus 41 µm long (33.6% of pharynx length) and 45 µm wide. Cephalic capsule 28 µm long, bacterial coat thick, consisting of coccobacilli approximately 1 µm long.

#### *Paralaxus* sp. “oahu 1”

(Figs. 43 – 45)

#### Sample locations

Oahu, Hawaii Islands, USA (21° 28’ 11,3’’ N, 157° 49’ 4,8’’ W and 21° 16’ 50,4’’ N, 157° 43’ 40,8 W)

#### Material

2 individuals, at least 1 male.

**Description** (measurements taken from LM micrographs of sequenced specimens)

Length 4927 µm

a= 71.4; b= 37.3; c= 35.7

Length of pharynx 132 µm, length of tail 138 µm; maximum diameter 69 µm, head diameter 41 µm, diameter at end of pharynx 58 µm, cloacal diameter 62 µm, c’= 2.2. Cuticle transversely striated, except for the cephalic capsule (34 µm long) and the tip of the tail, striae 1 µm wide, head and body setation not discernible in micrographs. Amphids in extreme forward position, open spiral with one turn, 13.5 µm wide, 9 µm long. Pharynx bulbus 36.4 µm wide and 38.8 µm long (29.6% of pharynx length). Spicula cephalate, curved, 64 µm (arc), 58 µm (chord) long, gubernaculum straight with terminal hook, 34.5 µm long. Several layers of coccobacilli approximately 1 µm long covering body.

## SI Tables

**Table S1:**
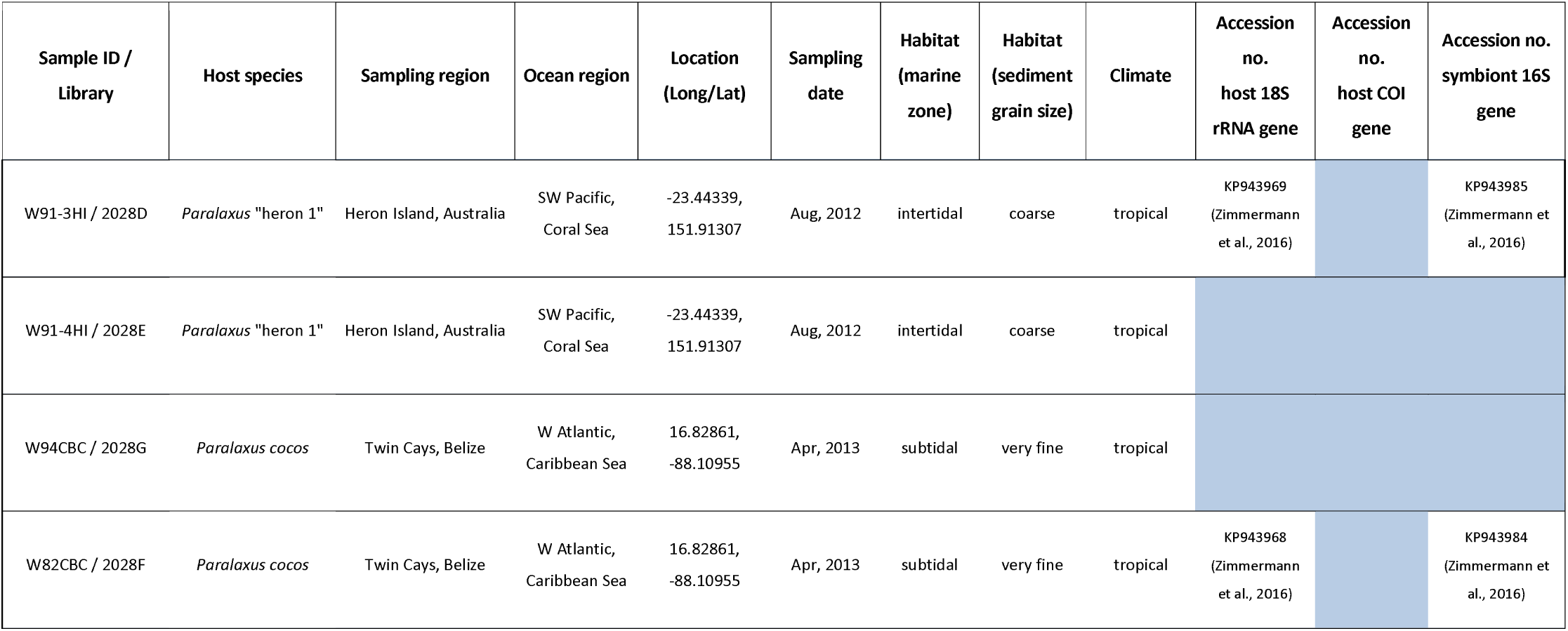
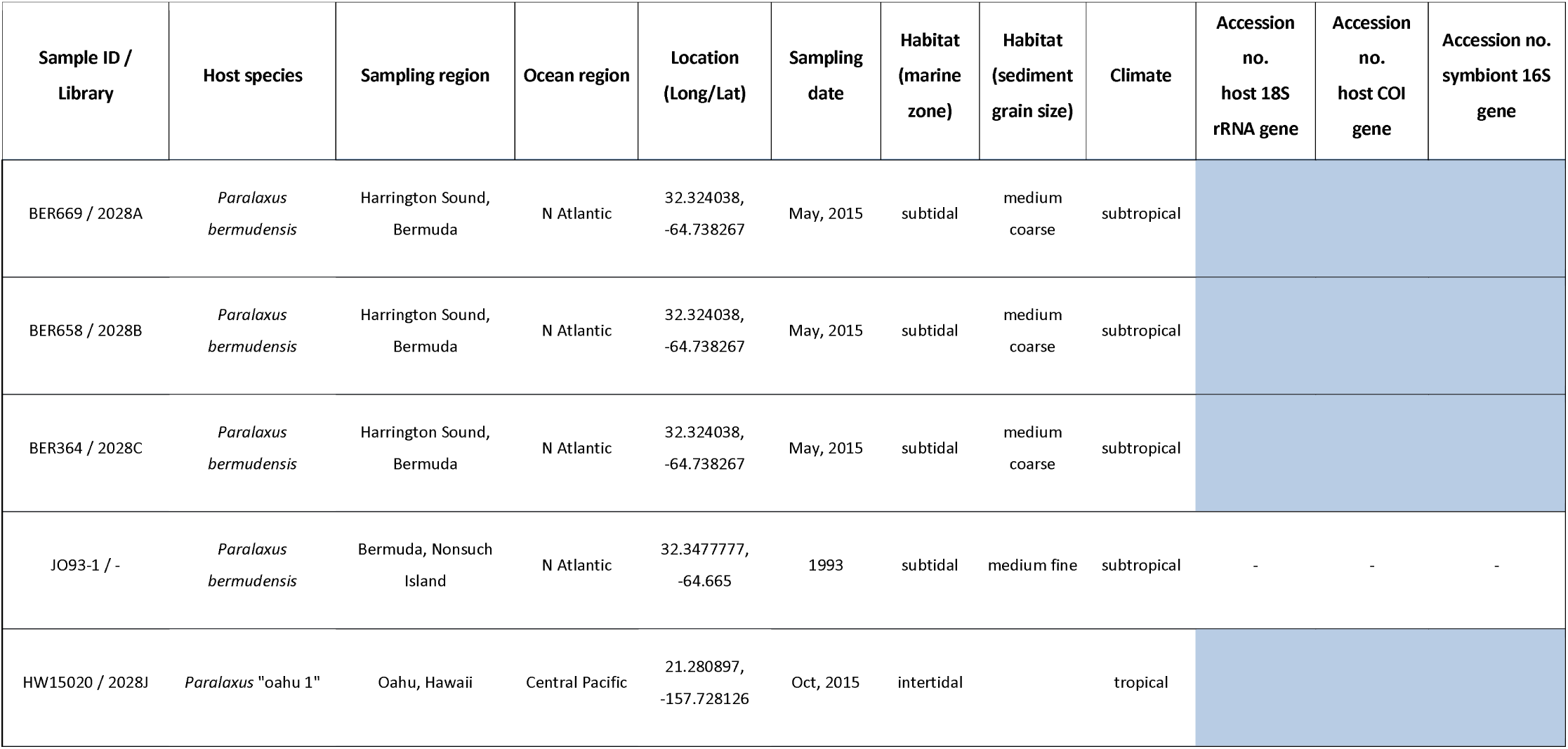
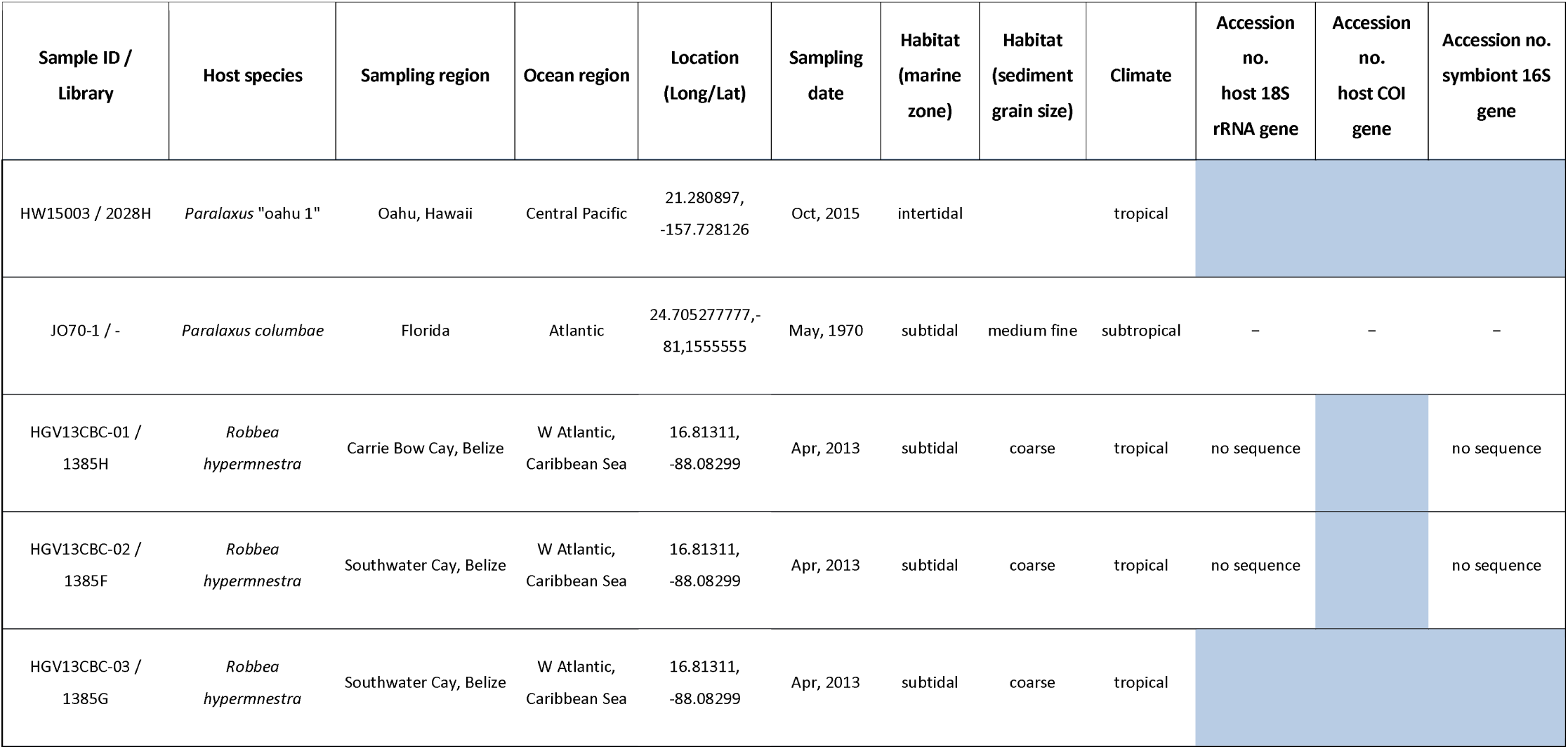
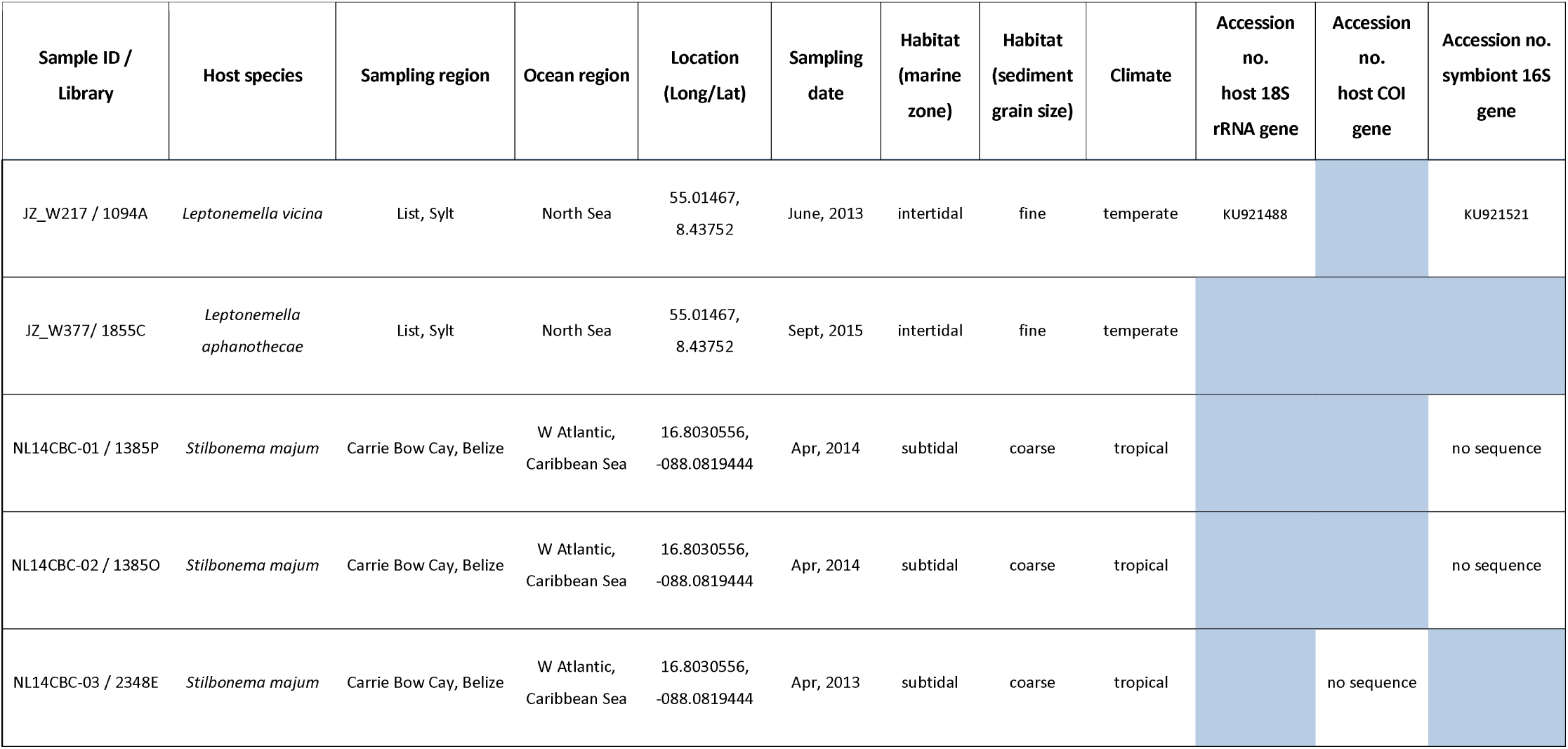
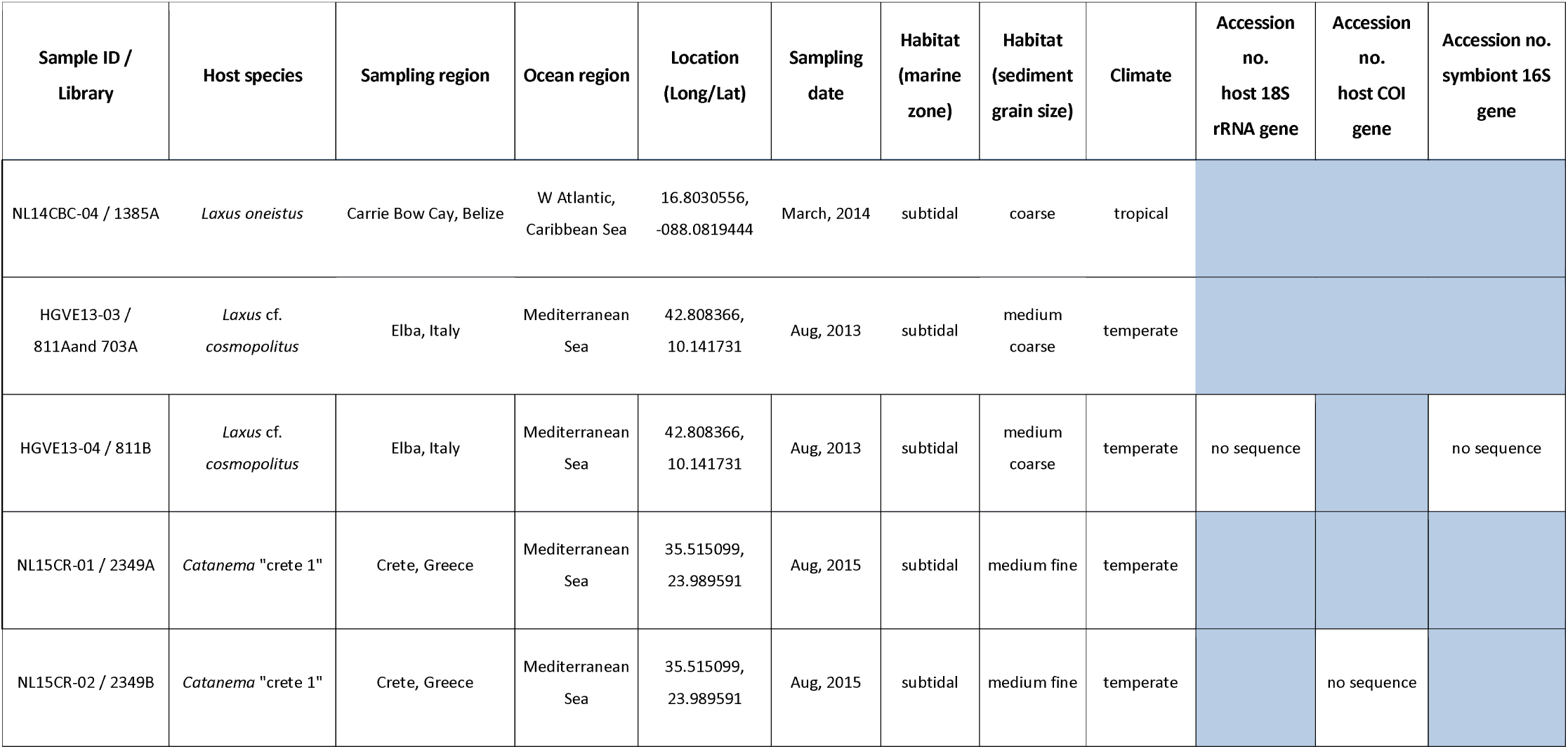
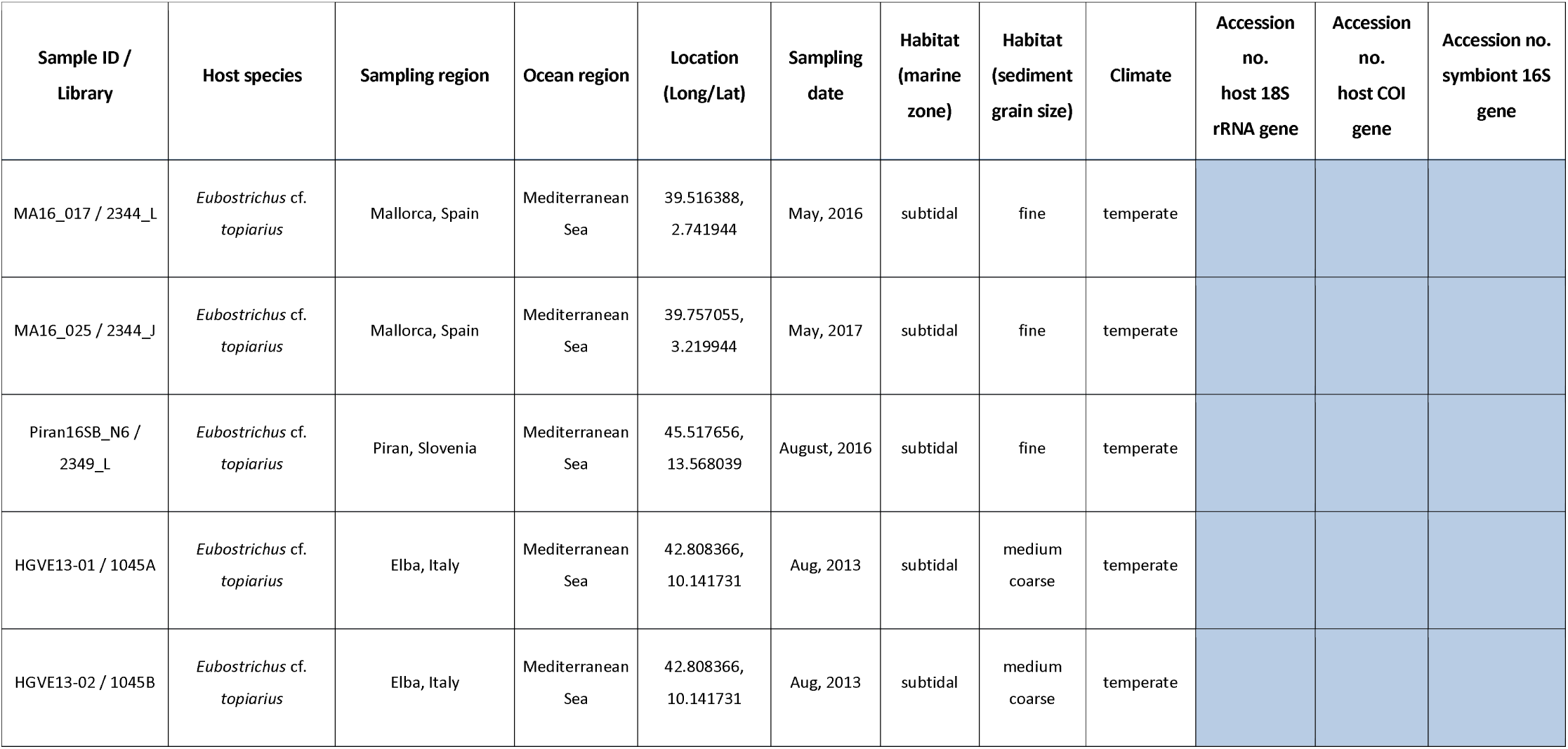
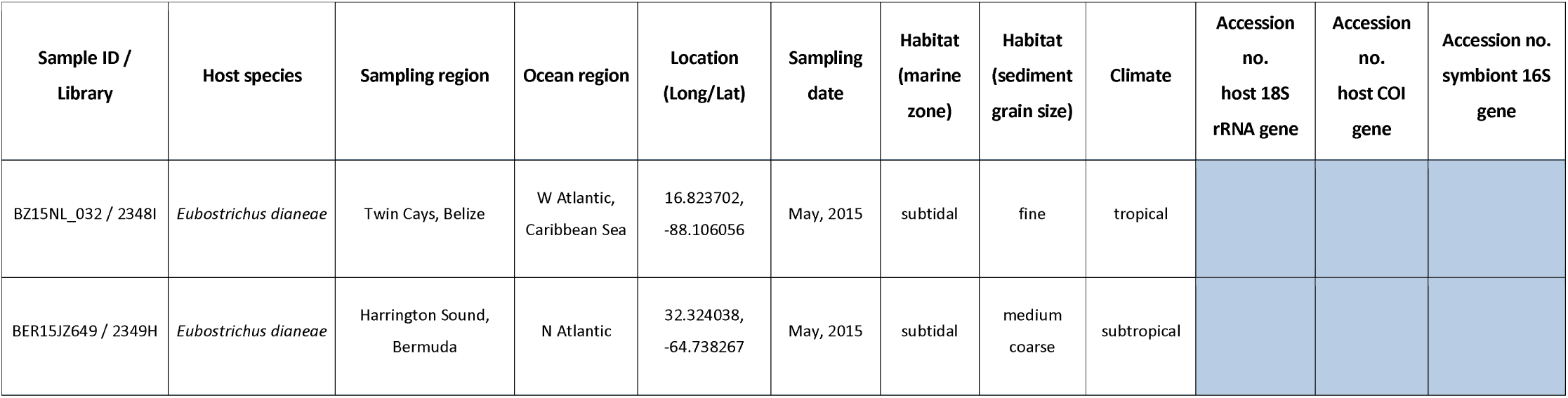
Stilbonematine nematode specimens sampled for this study.

| Sample ID / Library | Host species | Sampling region | Ocean region | Location (Long/Lat) | Sampling date | Habitat (marine zone) | Habitat (sediment grain size) | Climate | Accession no. host 18S rRNA gene | Accession no. host COI gene | Accession no. symbiont 16S gene |
| --- | --- | --- | --- | --- | --- | --- | --- | --- | --- | --- | --- |
| W91-3HI / 2028D | *Paralaxus "heron 1"* | Heron Island, Australia | SW Pacific, Coral Sea | -23.44339, 151.91307 | Aug, 2012 | intertidal | coarse | tropical | KP943969 (Zimmermann et al., 2016) |  | KP943985 (Zimmermann et al., 2016) |
| W91-4HI / 2028E | *Paralaxus "heron 1"* | Heron Island, Australia | SW Pacific, Coral Sea | -23.44339, 151.91307 | Aug, 2012 | intertidal | coarse | tropical |  |  |  |
| W94CBC / 2028G | *Paralaxus cocos* | Twin Cays, Belize | W Atlantic, Caribbean Sea | 16.82861, -88.10955 | Apr, 2013 | subtidal | very fine | tropical |  |  |  |
| W82CBC / 2028F | *Paralaxus cocos* | Twin Cays, Belize | W Atlantic, Caribbean Sea | 16.82861, -88.10955 | Apr, 2013 | subtidal | very fine | tropical | KP943968 (Zimmermann et al., 2016) |  | KP943984 (Zimmermann et al., 2016) |
| BER669 / 2028A | *Paralaxus bermudensis* | Harrington Sound, Bermuda | N Atlantic | 32.324038, -64.738267 | May, 2015 | subtidal | medium coarse | subtropical |  |  |  |
| BER658 / 2028B | *Paralaxus bermudensis* | Harrington Sound, Bermuda | N Atlantic | 32.324038, -64.738267 | May, 2015 | subtidal | medium coarse | subtropical |  |  |  |
| BER364 / 2028C | *Paralaxus bermudensis* | Harrington Sound, Bermuda | N Atlantic | 32.324038, -64.738267 | May, 2015 | subtidal | medium coarse | subtropical |  |  |  |
| JO93-1 / - | *Paralaxus bermudensis* | Bermuda, Nonsuch Island | N Atlantic | 32.3477777, -64.665 | 1993 | subtidal | medium fine | subtropical | - | - | - |
| HW15020 / 2028J | *Paralaxus* "oahu 1" | Oahu, Hawaii | Central Pacific | 21.280897, -157.728126 | Oct, 2015 | intertidal |  | tropical |  |  |  |
| HW15003 / 2028H | *Paralaxus* "oahu 1" | Oahu, Hawaii | Central Pacific | 21.280897, -157.728126 | Oct, 2015 | intertidal |  | tropical |  |  |  |
| JO70-1 / - | *Paralaxus columbae* | Florida | Atlantic | 24.705277777, -81.1555555 | May, 1970 | subtidal | medium fine | subtropical | - | - | - |
| HGV13CBC-01 / 1385H | *Robbea hypermnestra* | Carrie Bow Cay, Belize | W Atlantic, Caribbean Sea | 16.81311, -88.08299 | Apr, 2013 | subtidal | coarse | tropical | no sequence |  | no sequence |
| HGV13CBC-02 / 1385F | *Robbea hypermnestra* | Southwater Cay, Belize | W Atlantic, Caribbean Sea | 16.81311, -88.08299 | Apr, 2013 | subtidal | coarse | tropical | no sequence |  | no sequence |
| HGV13CBC-03 / 1385G | *Robbea hypermnestra* | Southwater Cay, Belize | W Atlantic, Caribbean Sea | 16.81311, -88.08299 | Apr, 2013 | subtidal | coarse | tropical |  |  |  |
| JZ_W217 / 1094A | *Leptonemella vicina* | List, Sylt | North Sea | 55.01467, 8.43752 | June, 2013 | intertidal | fine | temperate | KU921488 |  | KU921521 |
| JZ_W377/ 1855C | *Leptonemella aphanothecae* | List, Sylt | North Sea | 55.01467, 8.43752 | Sept, 2015 | intertidal | fine | temperate |  |  |  |
| NL14CBC-01 / 1385P | *Stilbonema majum* | Carrie Bow Cay, Belize | W Atlantic, Caribbean Sea | 16.8030556, -088.0819444 | Apr, 2014 | subtidal | coarse | tropical |  |  | no sequence |
| NL14CBC-02 / 1385O | *Stilbonema majum* | Carrie Bow Cay, Belize | W Atlantic, Caribbean Sea | 16.8030556, -088.0819444 | Apr, 2014 | subtidal | coarse | tropical |  |  | no sequence |
| NL14CBC-03 / 2348E | *Stilbonema majum* | Carrie Bow Cay, Belize | W Atlantic, Caribbean Sea | 16.8030556, -088.0819444 | Apr, 2013 | subtidal | coarse | tropical |  | no sequence |  |
| NL14CBC-04 / 1385A | *Laxus oneistus* | Carrie Bow Cay, Belize | W Atlantic, Caribbean Sea | 16.8030556, -088.0819444 | March, 2014 | subtidal | coarse | tropical |  |  |  |
| HGVE13-03 / 811Aand 703A | *Laxus cf. cosmopolitus* | Elba, Italy | Mediterranean Sea | 42.808366, 10.141731 | Aug, 2013 | subtidal | medium coarse | temperate |  |  |  |
| HGVE13-04 / 811B | *Laxus cf. cosmopolitus* | Elba, Italy | Mediterranean Sea | 42.808366, 10.141731 | Aug, 2013 | subtidal | medium coarse | temperate | no sequence |  | no sequence |
| NL15CR-01 / 2349A | *Catanema "crete 1"* | Crete, Greece | Mediterranean Sea | 35.515099, 23.989591 | Aug, 2015 | subtidal | medium fine | temperate |  |  |  |
| NL15CR-02 / 2349B | *Catanema "crete 1"* | Crete, Greece | Mediterranean Sea | 35.515099, 23.989591 | Aug, 2015 | subtidal | medium fine | temperate |  |  |  |
| MA16_017 / 2344_L | *Eubostrichus cf. topiarius* | Mallorca, Spain | Mediterranean Sea | 39.516388, 2.741944 | May, 2016 | subtidal | fine | temperate |  |  |  |
| MA16_025 / 2344_J | *Eubostrichus cf. topiarius* | Mallorca, Spain | Mediterranean Sea | 39.757055, 3.219944 | May, 2017 | subtidal | fine | temperate |  |  |  |
| Piran16SB_N6 / 2349_L | *Eubostrichus cf. topiarius* | Piran, Slovenia | Mediterranean Sea | 45.517656, 13.568039 | August, 2016 | subtidal | fine | temperate |  |  |  |
| HGVE13-01 / 1045A | *Eubostrichus cf. topiarius* | Elba, Italy | Mediterranean Sea | 42.808366, 10.141731 | Aug, 2013 | subtidal | medium coarse | temperate |  |  |  |
| HGVE13-02 / 1045B | *Eubostrichus cf. topiarius* | Elba, Italy | Mediterranean Sea | 42.808366, 10.141731 | Aug, 2013 | subtidal | medium coarse | temperate |  |  |  |

| Sample ID /<br>Library | Host species | Sampling region | Ocean region | Location<br>(Long/Lat) | Sampling<br>date | Habitat<br>(marine<br>zone) | Habitat<br>(sediment<br>grain size) | Climate | Accession<br>no.<br>host 18S<br>rRNA gene | Accession<br>no.<br>host COI<br>gene | Accession no.<br>symbiont 16S<br>gene |
| --- | --- | --- | --- | --- | --- | --- | --- | --- | --- | --- | --- |
| BZ15NL_032 / 2348I | *Eubostrichus dianeae* | Twin Cays, Belize | W Atlantic,<br>Caribbean Sea | 16.823702,<br>-88.106056 | May, 2015 | subtidal | fine | tropical |  |  |  |
| BER15JZ649 / 2349H | *Eubostrichus dianeae* | Harrington Sound,<br>Bermuda | N Atlantic | 32.324038,<br>-64.738267 | May, 2015 | subtidal | medium<br>coarse | subtropical |  |  |  |

**Table S2.** Morphological details and measurements taken from microphotographs of live specimens of *Paralaxus cocos, P. bermudensis* and *P. columbae.*

|  | *Paralaxus cocos* |  |  | *Paralaxus bermudensis* |  | *Paralaxus columbae* |  |  |
| --- | --- | --- | --- | --- | --- | --- | --- | --- |
|  | Holotype | Paratype male | Paratype female | Holotype | Paratype male | Holotype | Paratype male | Paratype female |
| n | 1 | 3 | 3 | 1 | 1 | 1 | 1 | 3 |
| Length | 3822 | 3135 -4602 | 4336 - 5069 | 4115 | 4336 | 3568 | 2801 | 3935 - 4669 |
| a | 73.5 | 56.9 - 83.6 | 72.3 - 94.3 | 68.6 | 66.7 | 64.9 | 50.9 | 66 - 78 |
| b | 42.9 | 28.5 - 43 | 36.7 - 43.4 | 35.5 | 34.7 | 32.4 | 28 | 35.8 - 44.5 |
| c | 31.8 | 27.3 - 41.7 | 40.8 - 42.9 | 39.2 | 34.7 | 31.8 | 25.9 | 36.4 - 39.7 |
| max. diameter | 52 | 50 - 55 | 46 - 66 | 60 | 65 | 55 | 55 | 55 - 60 |
| pharynx length | 89 | 98 - 110 | 100 - 130 | 116 | 125 | 110 | 100 | 103 - 110 |
| tail length | 120 | 108 - 115 | 102 - 118 | 105 | 125 | 112 | 108 | 100 - 120 |
| anal/cloacal diameter | 50 | 50 - 55 | 40 - 42 | 60 | 65 | 55 | 52 | 50 - 55 |
|  | Holotype | Paratype male | Paratype female | Holotype | Paratype male | Holotype | Paratype male | Paratype female |
| c' | 2.4 | 2.0 - 2.3 | 2.6 - 2.8 | 1.75 | 1.9 | 2 | 1.96 | 1.8 - 2.2 |
| vulva % | n.a. | n.a. | 44 - 45 | n.a. | n.a. | n.a. | n.a. | 44.5 - 48 |
| amphidial fovea width | 14 | 14 - 15 | 11 bis 12 | 15 | 12 | 12 | 12 | 12 bis 13 |
| pharynx bulbus l/w | 30/28 | 32-37/32-35 | 30-45/27-43 | 37/35 | 42/40 | 35/32 | 30/35 | 30-35/37-45 |
| bulbus % of pharynx l | 33.7 | 31.8 - 34.5 | 30 - 34.6 | 31.9 | 32 | 31.8 | 30 | 29 - 33 |
| diam. end of pharynx | 47 | 46 - 57 | 46 - 50 | 60 | 65 | 55 | 54 | 55 - 60 |
| spicula length, arc/chord | 61/50 | 60-65/50-51 | n.a. | 57/45 | 63/45 | 60/50 | 60/50 | n.a. |
| gubernaculum length | 33 | 31 - 35 | n.a. | 30 | 30 | 30 | 32 | n.a. |
| velum % of tail length | 47 | 43 - 46 | n.a. | 40 | 36 | 35.7 | 37 | n.a. |

## SI Figures

**Fig. S1.**
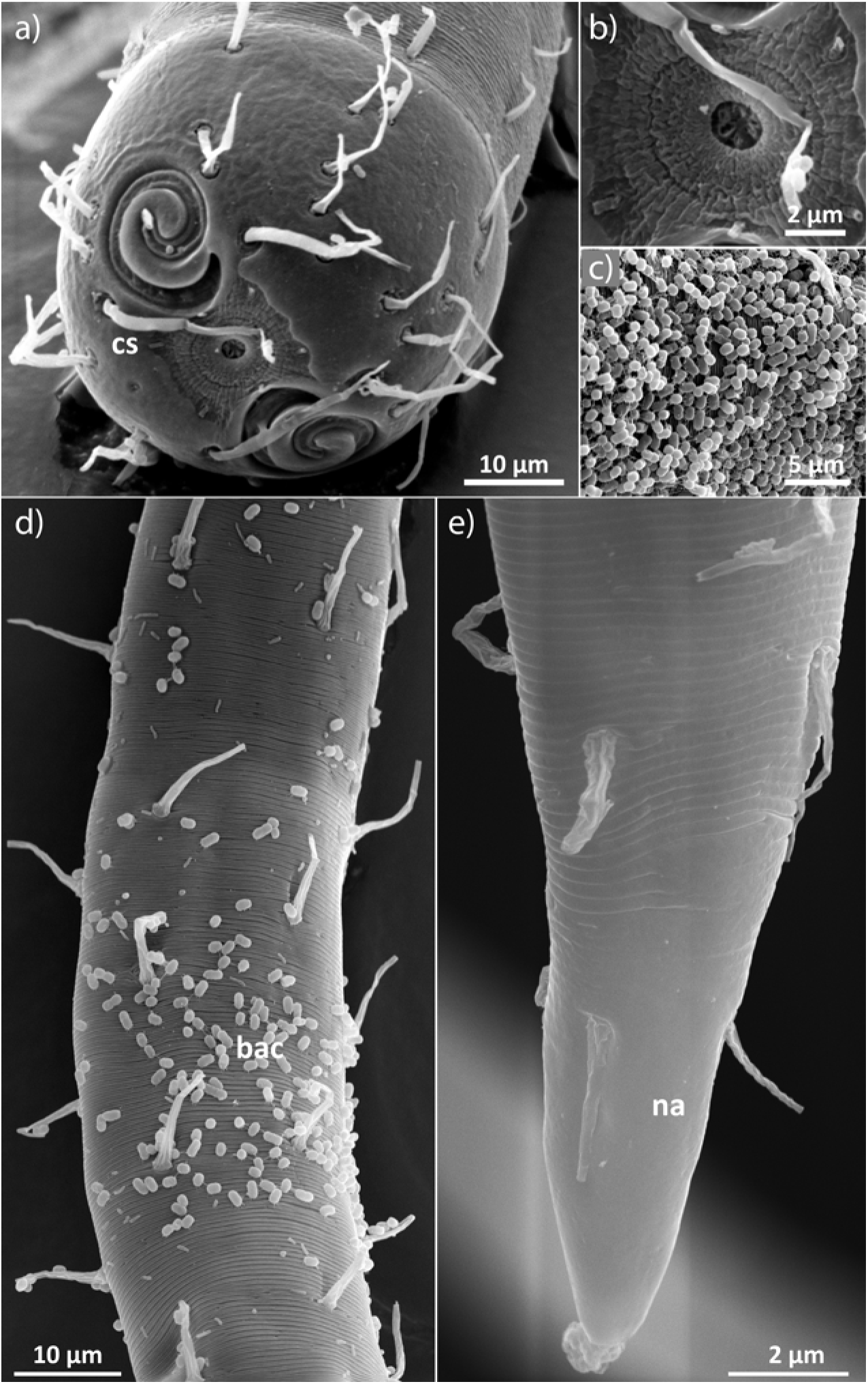
Scanning electron micrographs of *Paralaxus cocos* nov. spec.female. (a) Anterior in *en face* view, (b) buccal field, (c) detail of symbiotic coat, (d) midbody region, (e) tip of tail. SEM (ann annulation, bac symbiotic bacteria, bf buccal field, cs cephalic setae, fa fovea amphidalis, ls labial setae, na non–annulated tip of tail, scs subcephalic setae, ss somatic setae.

**Fig. S2.**
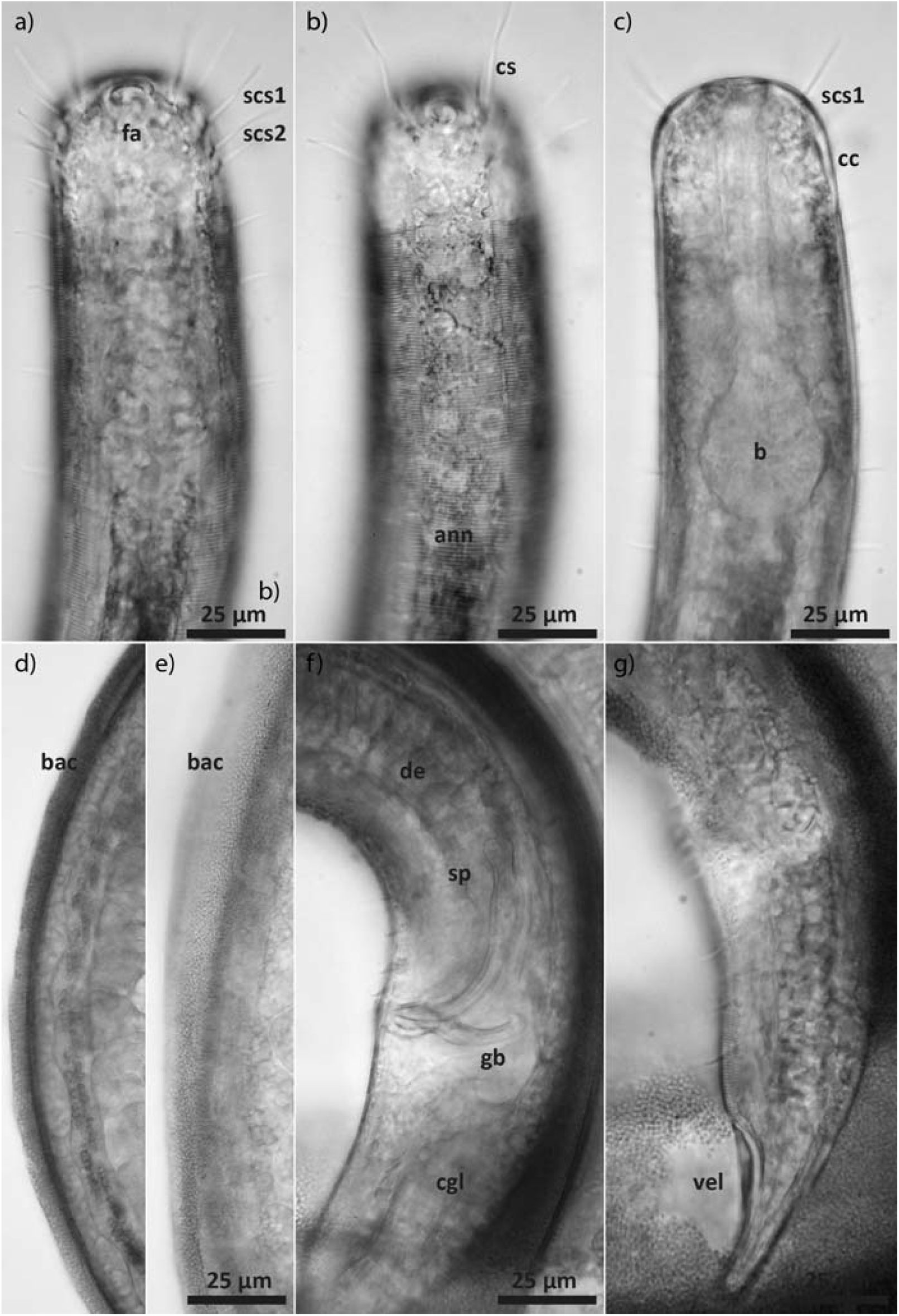
Light micrographs of *Paralaxus cocos* nov. spec. male. (a) anterior end (b) anterior end surface view (c) anterior end, optical section (d) midbody region, optical section of bacterial coat (e) midbody region, surface view of bacterial coat (f) cloacal region (g) tip of tail. LM of live specimen. (ann annulation, b pharyngeal bulbus, bac symbiotic bacteria, cc cephalic capsule, cgl caudal gland, cs cephalic setae, d ductus ejaculatorius, fa fovea amphidalis, gb gubernaculum, scs1 subcephalic setae of first circle, scs2 subcephalic setae of second circle, sp spiculum, vel velum).

**Fig. S3.**
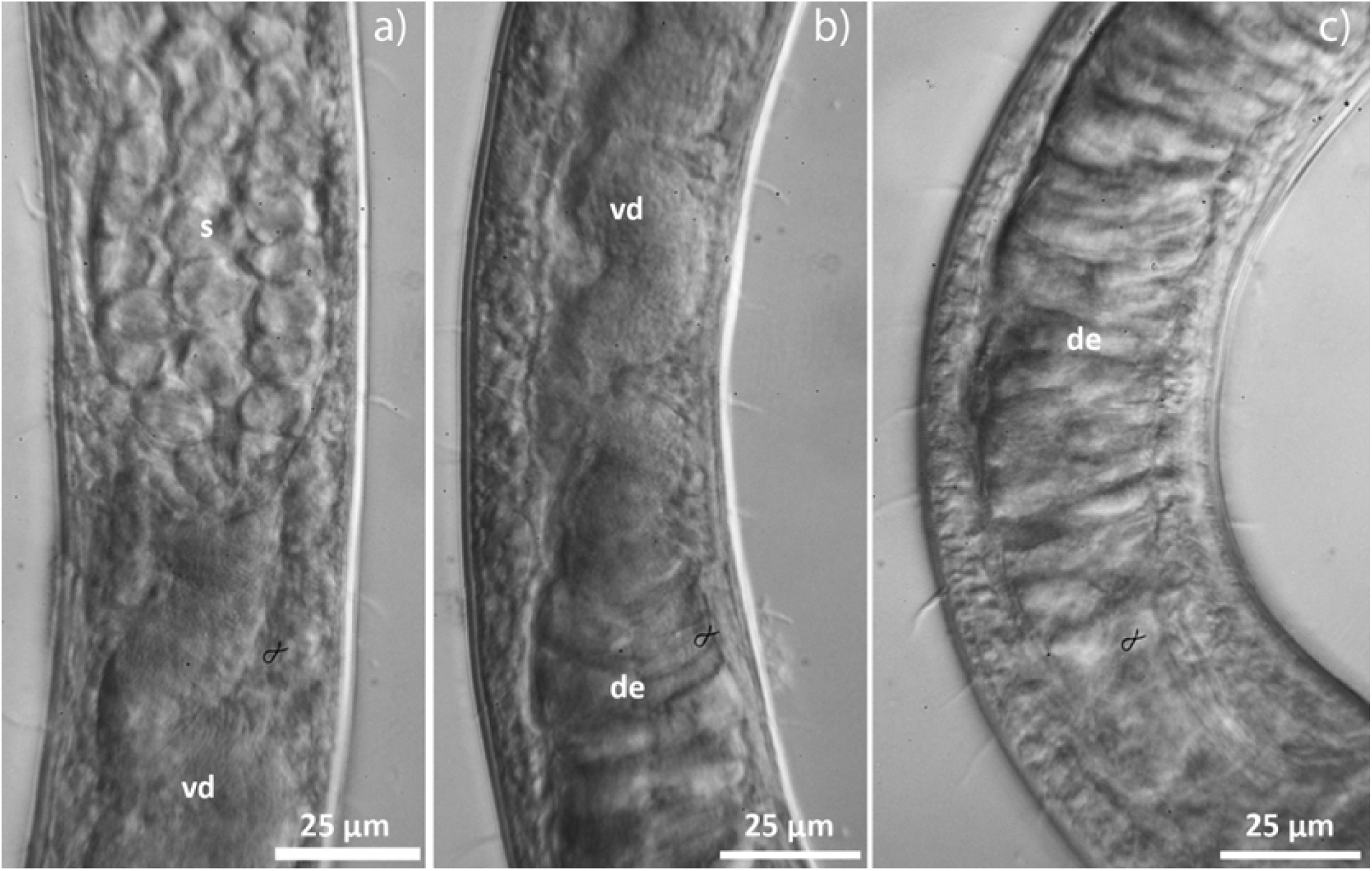
Light micrographs of the reproductive system of *Paralaxus cocos* nov. spec. male. (a) junction of testis and vas deferens (b) junction of vas deferens and ductus ejaculatorius; (c) posterior end of ductus ejaculatorius. LM of live specimen. (de ductus ejaculatorius, s sperm, t testis, vd vas deferens)

**Fig. S4.**
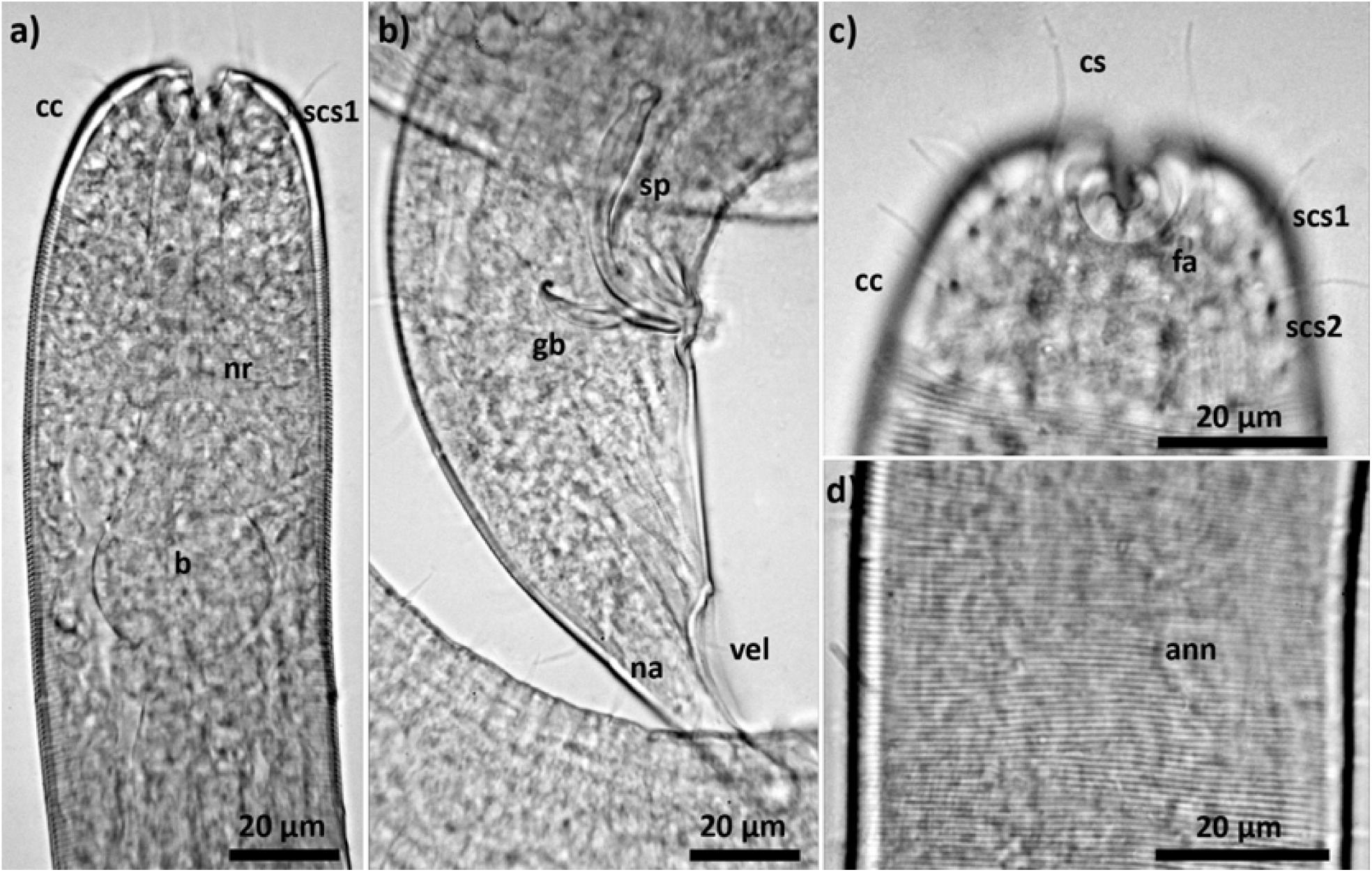
Light micrographs of *Paralaxus bermudensis* nov. spec. male. (a) anterior end, optical section (b) posterior end (c) anterior tip, surface view (d) midbody region. LM of preserved specimen. (ann annulation, b pharyngeal bulbus, cc cephalic capsule, cs cephalic setae, fa fovea amphidalis, gb gubernaculum, na non–annulated tip of tail, nr nerve ring, scs1 subcephalic setae of first circle, scs2 subcephalic setae of second circle, sp spiculum, vel velum).

**Fig. S5.**
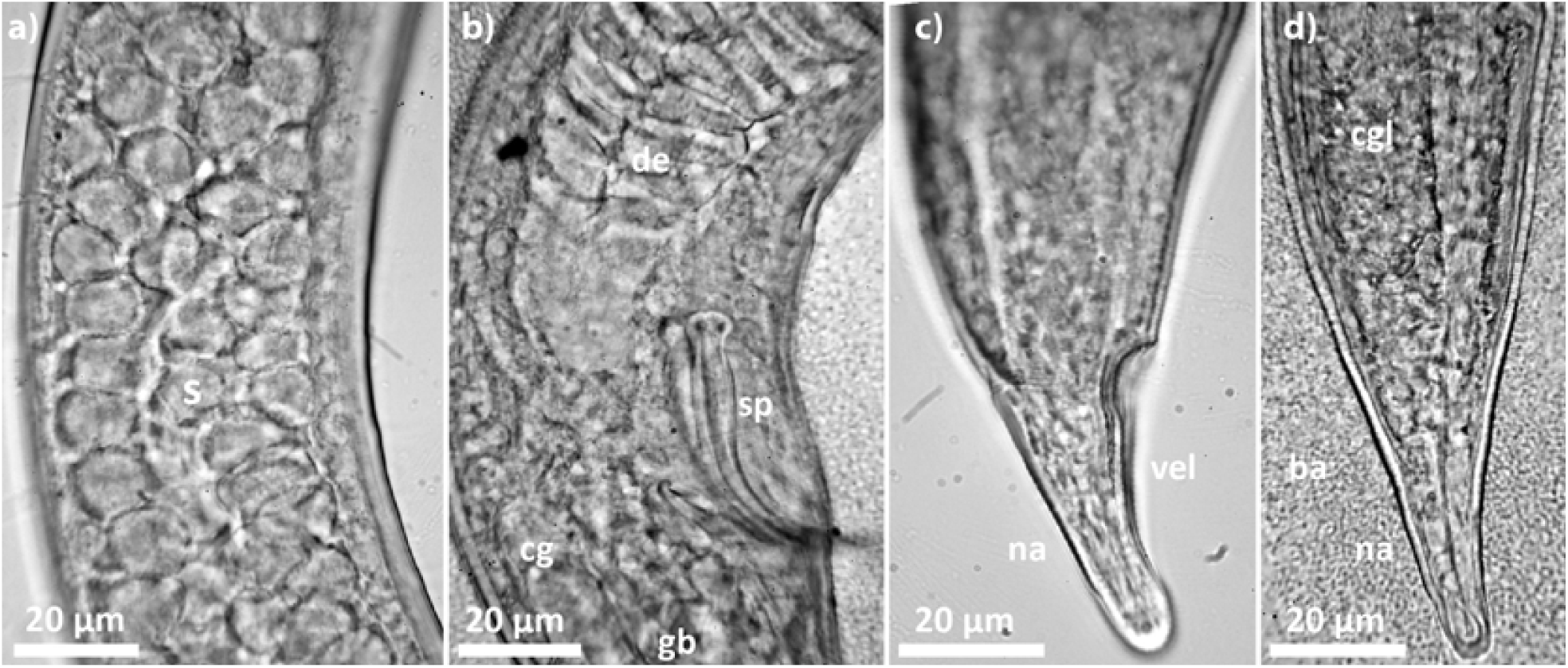
Light micrographs of the posterior end of *Paralaxus columbae* nov. spec. (a – c) male (a) sperm in testis (b) cloacal region (c) tail (d) female tail. LM of preserved specimen (bac symbiotic bacteria, cgl caudal gland, d ductus ejaculatorius, gb gubernaculum, na non– annulated tip of tail, sp spiculum, vel velum).

**Fig. S6.**
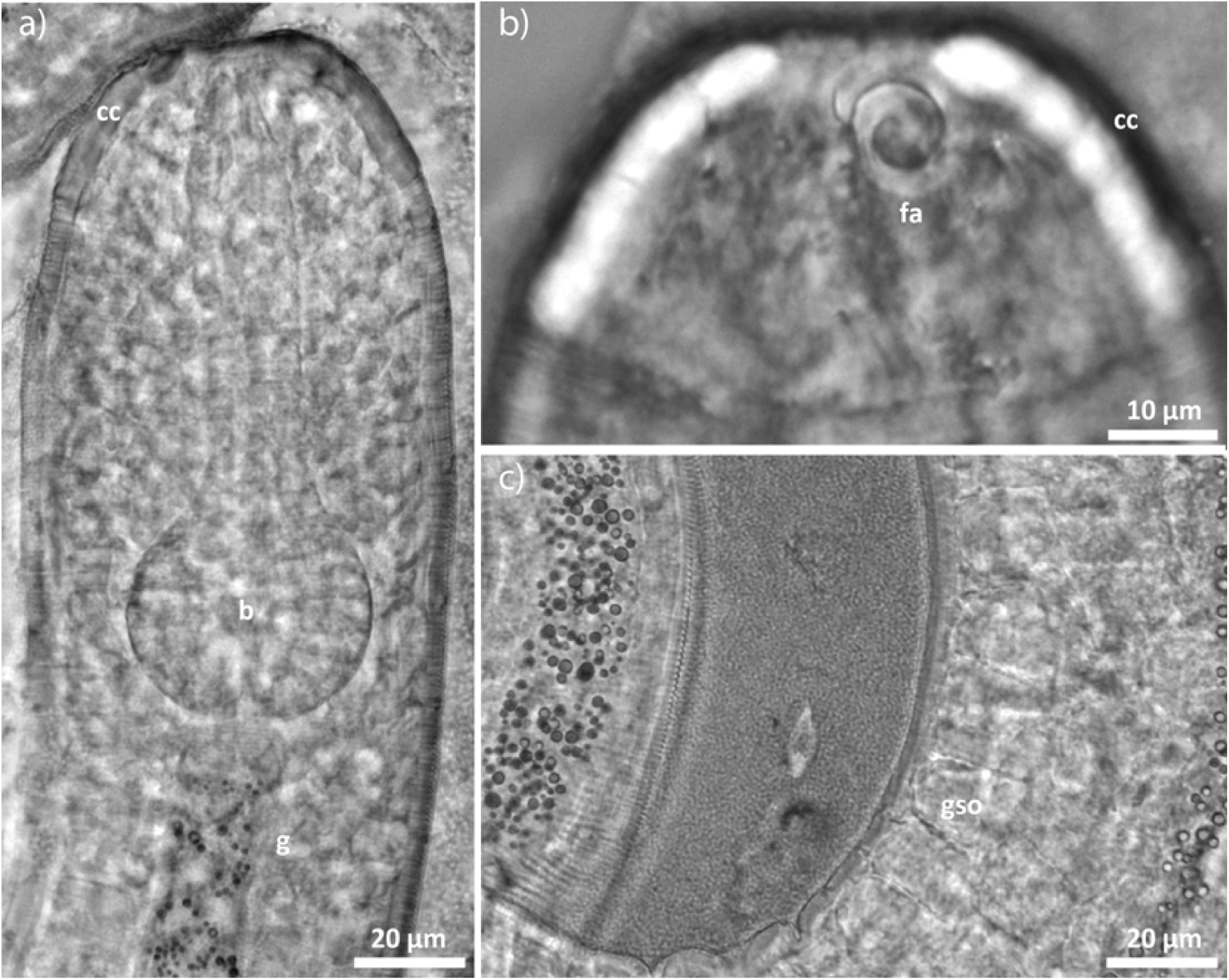
Light micrographs of *Paralaxus* sp. “heron”. (a) anterior region (b) anterior end (c) midbody region with dissociated bacterial coat. LM of preserved specimen. (b pharyngeal bulbus, bac bacteria, cc cephalic capsule, fa fovea amphidialis, g gut, gso glandular sense organ).

**Fig. S7.**
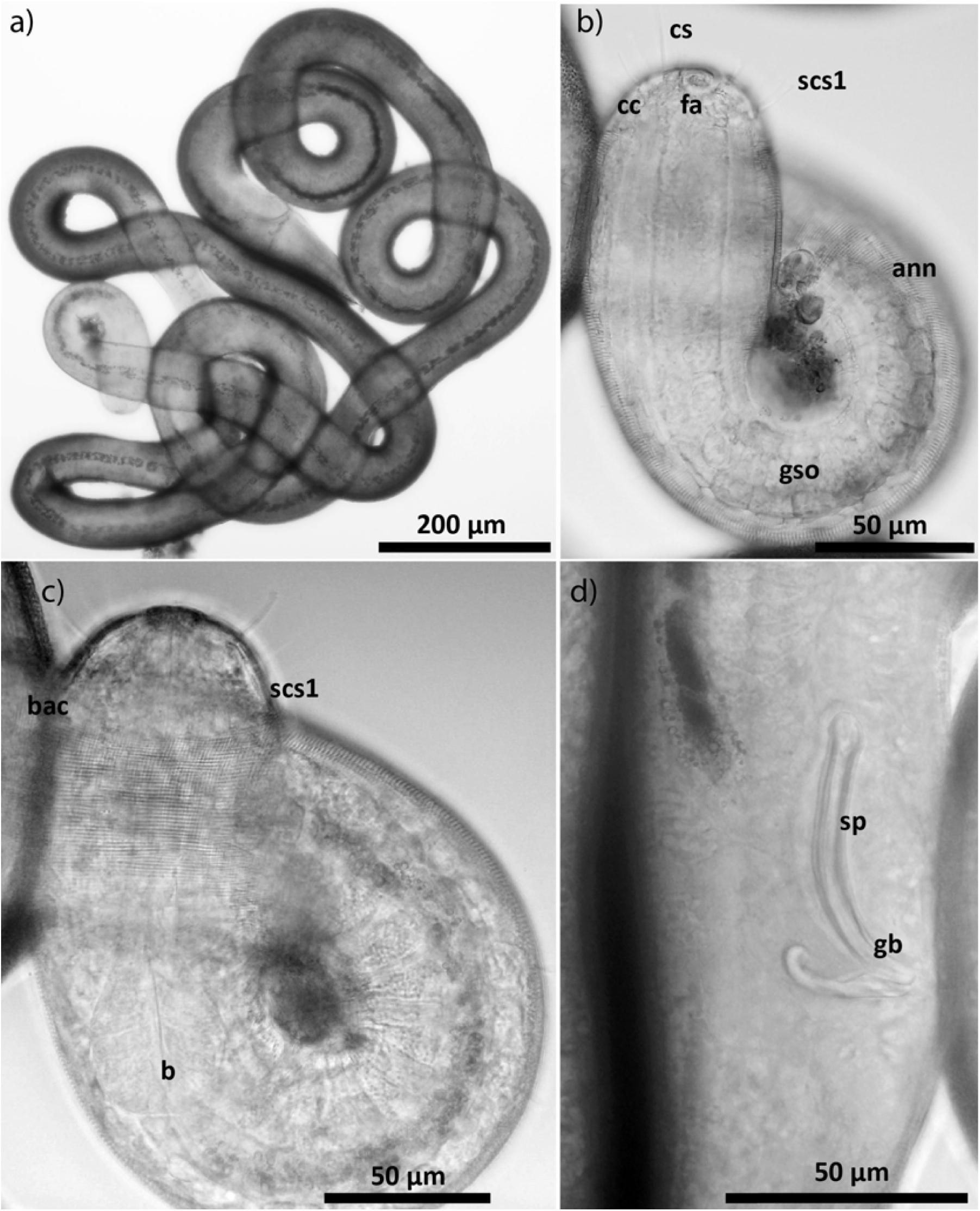
Light micrographs of *Paralaxus* “oahu” male. (a) total view (b) anterior region (c) optical section of anterior region (d) cloacal region. LM of live specimen (ann annulation, b pharyngeal bulbus, bac bacteria, cc cephalic capsule, cs cephalic setae, gb gubernaculum, gso glandular sense organ, scs1 subcephalic setae of first circle, sp spiculum).

**Fig. S8.**
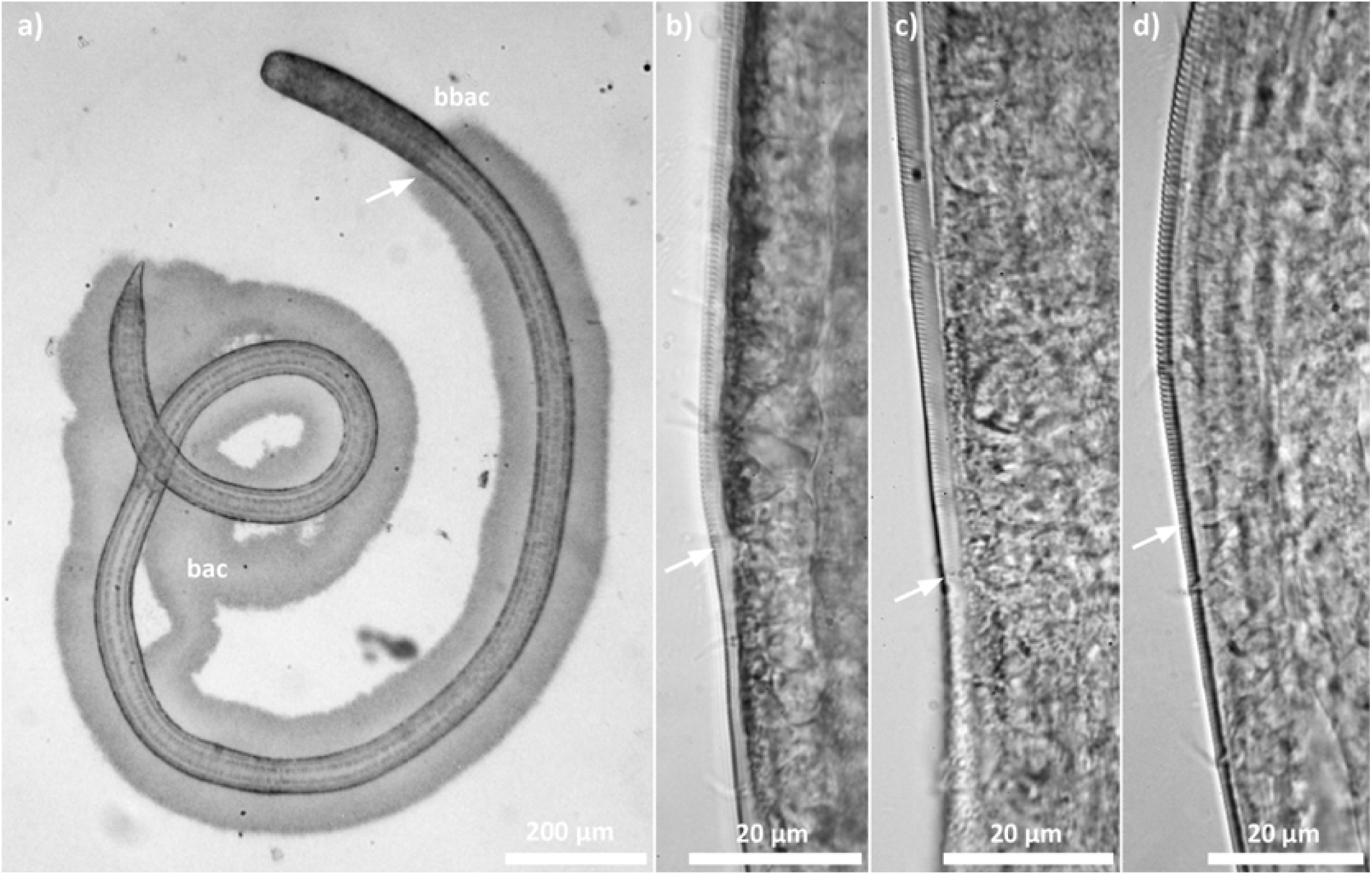
Light micrographs of *Paralaxus* species and its morphological adaptation to its symbiont coat. (a) *Paralaxus columbae* nov. spec, juvenile. Bacterial coat starting at the transition from coarse to finer annulation; (b – d) Transition in adult specimens (b) *P. cocos* nov. sp. (c) *P. columbae* nov. sp. (d) *P. bermudensis* nov. sp. The transition points are marked by white arrows. LM of preserved specimen. (bac bacteria, bbac begin of bacterial coat)

**Fig. S9.**
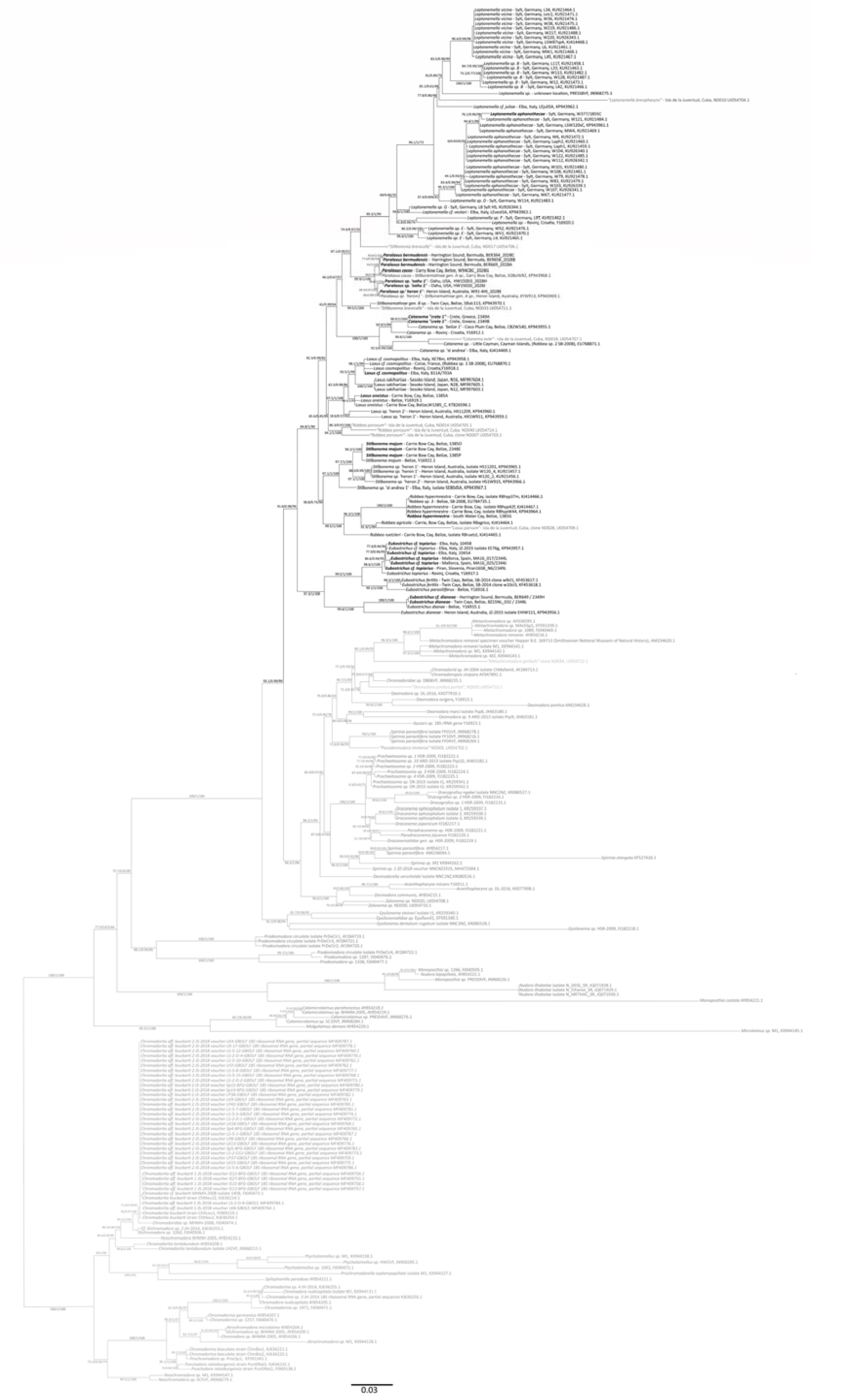
Host 18S rRNA phylogeny of extended dataset (non–collapsed tree version) The tree was calculated using IQ–Tree. Support values are given in the following order: SH–aLRT support (%) / aBayes support / ultrafast bootstrap support (%). Species described in this study are highlighted in bold. Species from Armenteros et al. (2014b) are highlighted in italic and light grey, with quote marks. Provisional working names for undescribed genera or species are given in quotes. The scale bar represents average nucleotide substitutions per site.

**Fig. S10.**
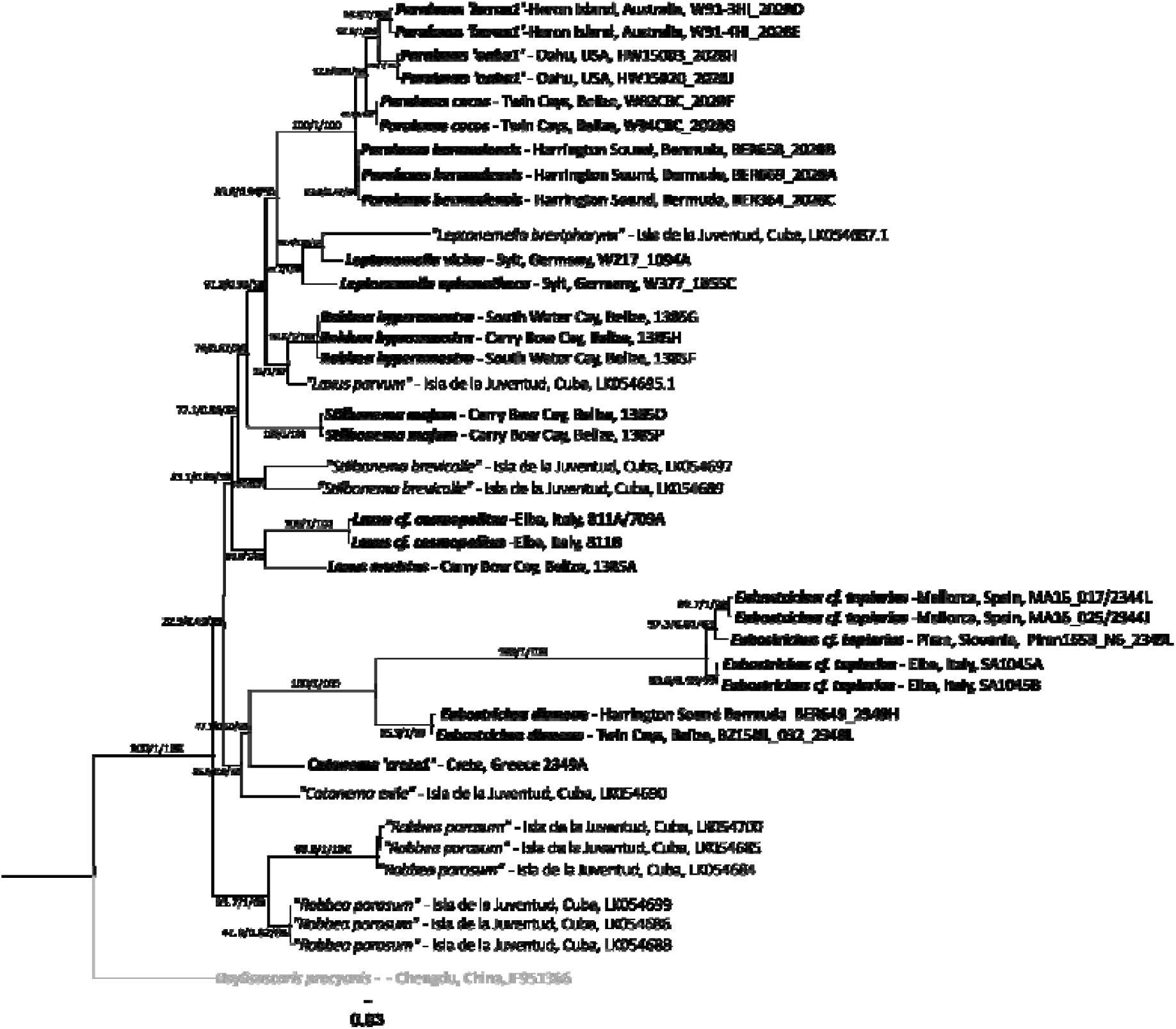
Host COI phylogeny of extended dataset (non–collapsed tree version) The tree was calculated using IQ–Tree. Support values are given in the following order: SH–aLRT support (%) / aBayes support / ultrafast bootstrap support (%). Species described in this study are highlighted in bold. Species from Armenteros et al. (2014b) are highlighted in italic and with quote marks. Outgroup in light grey and italic. Provisional working names for undescribed genera or species are given in quotes. The scale bar represents average nucleotide substitutions per site.

**Fig. S11.**
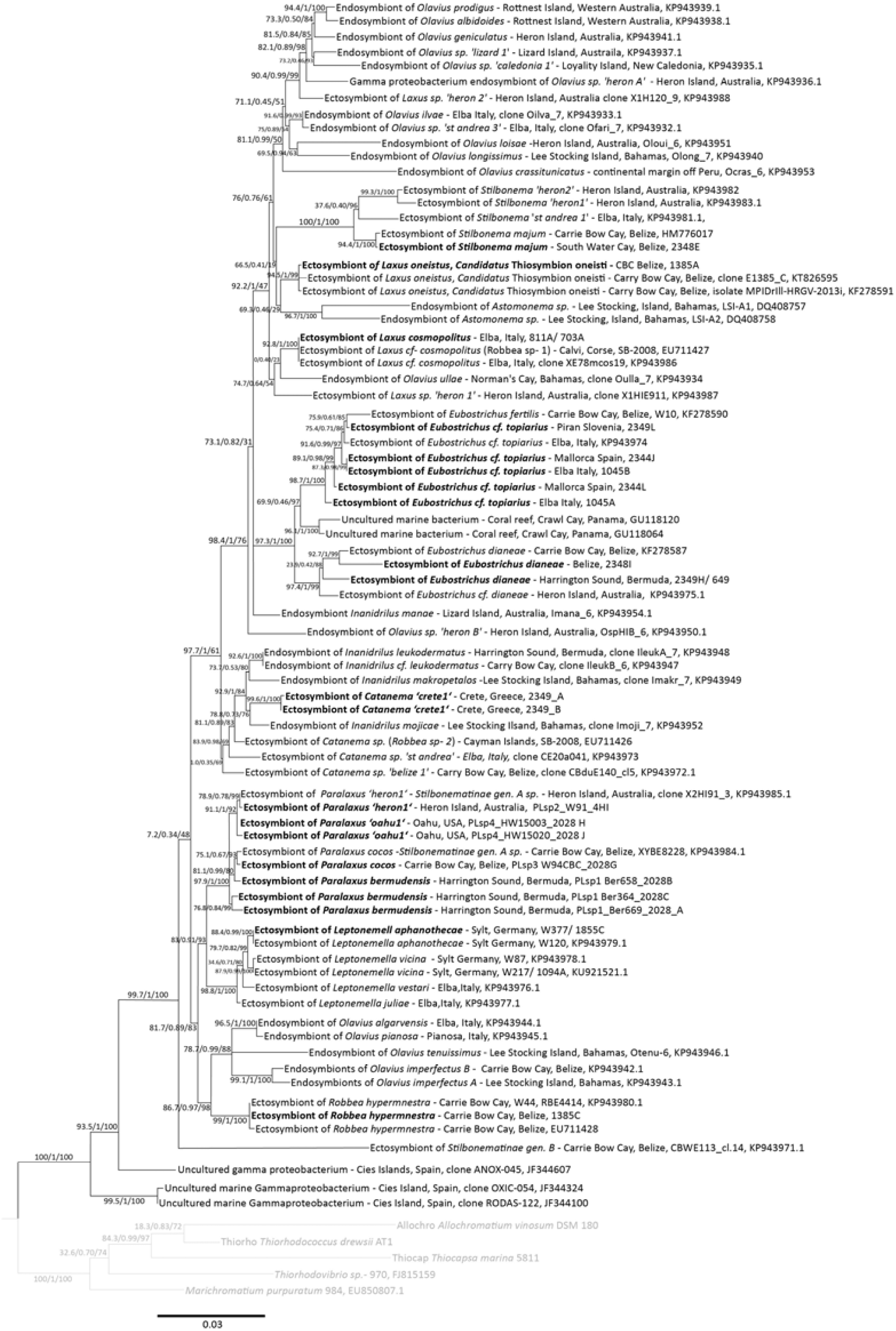
Symbiont 16S rRNA phylogeny (non–collapsed tree version) The tree was calculated using IQ–Tree. Support values are given in the following order: SH–aLRT support (%) / aBayes support / ultrafast bootstrap support (%). Symbiont sequences generated in this study are highlighted in bold. Outgroup in light grey. Provisional working names for undescribed genera or species are given in quotes. The scale bar represents average nucleotide substitutions per site.

